# Data mining of global sequence datasets markedly expands known diversity within the Tobamovirus genus

**DOI:** 10.64898/2026.10.22.760918

**Authors:** Neža Pajek Arambašič, Katarina Bačnik, Luka Kranjc, Lana Vogrinec, Tomaž Curk, Denis Kutnjak

## Abstract

High-throughput sequencing has transformed RNA virus discovery, producing vast datasets that often contain unexplored sequences of known and novel viruses across diverse sample types. Large-scale reconstruction of viral genomes within specific taxonomic groups remains a lengthy, labor-intensive process constrained by fragmented assemblies, heterogeneous workflows, and the limitations of percent identity cut-offs for taxonomic placement. Here, we focused on *Tobamovirus* (family *Virgaviridae*), a genus that contains important emerging plant pathogens. Tobamoviruses are detectable in diverse environmental matrices and are thus expected to be present across a range of datasets. We mined global short-read datasets to uncover hidden tobamovirus diversity, combining expert curation with supervised machine learning to streamline the discovery of novel tobamoviruses. Using the Serratus PalmID tool, we first identified 253 candidate datasets that are most likely to contain novel tobamoviruses. We developed a reproducible Snakemake pipeline for quality control, de novo assembly, similarity searches, and preliminary taxonomic assignment, yielding 510 candidate tobamovirus contigs across 131 datasets. Following manual curation, we confirmed 228 tobamoviral contigs from 23 datasets based on BLAST similarity searches, coding sequence content, genome organization, and assessment of potential misassemblies. To expedite the curation process, we built a two-stage machine learning classifier of tobamoviral contigs. Based on alignment-derived features and sequence properties, each ORF is first scored to determine whether it is tobamoviral. ORF-level predictions are then merged into contig-level classification using stacked generalization. Through clustering and phylogenetic analyses, we identified at least 32 putative novel tobamoviruses in datasets derived from freshwater, wastewater, plants, animal gut, and soil. Most were placed within established *Tobamovirus* subclades, whereas a few highly divergent sequences formed distinct basal branches related to recently identified soil-associated sequences classified as *Virgaviridae* sp. These findings highlight the value of aquatic and wastewater systems for expanding our understanding of tobamoviral diversity and demonstrate how data mining and targeted curation can substantially expand recognized diversity within the genus. The workflow is readily adaptable to other viral taxa and supports continuous discovery as public datasets grow.

## Introduction

In recent years, high-throughput sequencing-based metagenomic approaches have emerged as key methods in RNA virus discovery. Large-scale virome studies have significantly expanded our understanding of RNA viruses associated with organisms from different domains of life and environmental samples (Shi et al., 2016; Zayed et al., 2022; Rivarez et al., 2023). Shotgun sequencing data are generated across various life science fields and continually accumulate in public repositories. Even if not intended for virome analysis, such datasets hold valuable information on known and novel viruses, obtainable through re-analysis using data mining (Edgar et al., 2022; Neri et al., 2022; Mifsud et al., 2022; Vogrinec et al., 2025).

Several tools have been developed to facilitate data mining (Edgar et al., 2022; Chikhi et al., 2025; Karasikov et al., 2025) and exploration of repositories for viral sequences. For RNA viruses, the Serratus infrastructure (Edgar et al., 2022) provides an opportunity to explore all short-read datasets (from the NCBI Sequence Read Archive (SRA) database, up to February 2022) for the presence of highly conserved palmprint domains within the RNA-dependent RNA polymerase (RdRp), a hallmark of RNA viruses.

One promising application of data mining is the identification of previously unknown viruses that are phylogenetically related to known viral pathogens. Uncovering hidden viral diversity in the virosphere may enhance preparedness for future disease outbreaks. Viruses with stable virions that persist outside their hosts and are detectable across multiple sample types, and thus across diverse associated datasets, are good candidates for exploring this approach. Members of the genus *Tobamovirus* (family *Virgaviridae*), which includes the first virus ever discovered (*i*.*e*., tobacco mosaic virus, TMV), are among the most extensively studied plant viruses (Gibbs et al., 2015; Scholthof et al., 2011; Creager et al., 1999; Spiegelman and Dinesh-Kumar, 2023). Their virions are exceptionally stable, remaining infectious in the environment for extended periods (Mehle et al., 2018, 2023; Li et al., 2016). They have been detected in high abundance across various environmental matrices using both targeted (Rosario et al., 2009; Fillhart et al., 1998; Castello et al., 1995; Koenig, 1986) and metagenomic approaches (Bačnik et al., 2020; Rothman and Whiteson, 2022). In recent years, emerging tobamoviruses, such as tomato brown rugose fruit virus (ToBRFV), which can overcome resistance in commercial crop varieties, have caused substantial economic losses in agriculture (Caruso et al., 2022).

Despite extensive research on this genus and the agricultural importance of several of its members, putative novel tobamoviruses have recently been identified from both plant (Rivarez et al., 2023) and environmental samples (Maksimović et al., 2024). Given their environmental stability, we expect to find novel tobamoviruses in a wide range of sample types, including plants, water, wastewater, feces, and soil.

Tobamoviruses have a non-segmented, positive-sense single-stranded RNA genome of 6.3–6.6 kb, which codes for: two replication-associated proteins (124–132 kDa and 181–189 kDa), a movement protein (28–31 kDa), and a coat protein (17–18 kDa) (Black et al., 2026). As of July 2026, there are 37 recognized species within the *Tobamovirus* genus (Black et al., 2026). According to the criteria established by the International Committee on Taxonomy of Viruses (ICTV), species in the *Tobamovirus* genus are distinguished by 90% whole-genome nucleotide sequence identity, antigenic relationships among coat proteins, and host range (Black et al., 2026).

Species demarcation for tobamoviruses is performed using pairwise nucleotide sequence identity across complete genomes, typically with tools such as the Sequence Demarcation Tool (SDT) (Muhire et al., 2014). This approach is effective when full-length genomes are available. However, when mining public sequencing repositories, recovered viral sequences are often short, fragmented contigs due to low viral abundance, suboptimal sequencing depth, or an experimental design not tailored for virome analysis. A large number of these partial sequences pose significant challenges for accurate, time-efficient retrieval and classification of novel viral sequences based on pairwise sequence identities, including the need to establish the correct sequence orientation prior to analysis with SDT (Muhire et al., 2014).

The scale of data mining efforts adds another layer of complexity. While initial steps, such as identifying promising datasets, can be efficiently addressed using tools like the Serratus (Edgar et al., 2022), analyzing hundreds or more datasets to identify, assemble, and classify tobamoviral sequences remains computationally intensive and time-consuming. Each dataset must be processed individually, and candidate viral contigs should ideally be carefully curated to minimize mis-classifications. Manual curation of large-scale viral discovery datasets is labor-intensive and time-consuming. This challenge is exacerbated because assembled contigs often represent non-overlapping genome fragments and may contain only partial protein-coding sequences. Moreover, pairwise alignment tools can return inconsistent classifications when different algorithms are used (Bao et al., 2014), and sequence identity alone is often insufficient for classifying either closely related (Gibbs and Gibbs, 2018) or highly divergent taxa (Simmonds, 2015). Additional alignment-derived metrics, such as alignment length and gap ratio, can provide important context for taxonomic assignment but are not directly used in traditional cut-off-based workflows. These challenges underscore the need for scalable, accurate, and virus-specific bioinformatic pipelines capable of processing large volumes of metagenomic data to target the discovery of specific viral taxa.

In this study, we aimed to expand our understanding of the hidden diversity of tobamoviruses by mining publicly available sequencing datasets. To overcome the outlined challenges, we developed a modular workflow tailored for the discovery of tobamoviruses (see Figure 1). We first used the Serratus toolkit (Edgar et al., 2022) to identify the most promising short-read datasets. Next, we implemented an automated pipeline using Snakemake (v7.32.4) (Mölder et al., 2021), which included *de novo* assembly, similarity searches, and preliminary taxonomic classification to extract putative tobamoviral contigs (Pajek Arambašič et al., 2026b). We then pursued two complementary strategies: first, we manually curated and annotated the candidate tobamoviral contigs, including phylogenetic analyses of representative sequences. Second, we developed a machine learning model to streamline and partially automate downstream steps. This integrated approach enabled efficient data processing and characterization of tobamoviral sequences across large-scale metagenomic data. Ultimately, the workflow facilitated the discovery of at least 32 putative novel tobamoviruses and can be readily adapted to identify other viral taxa or to continue discovering tobamoviruses as new sequencing data become available in public repositories.

**Fig. 1:**
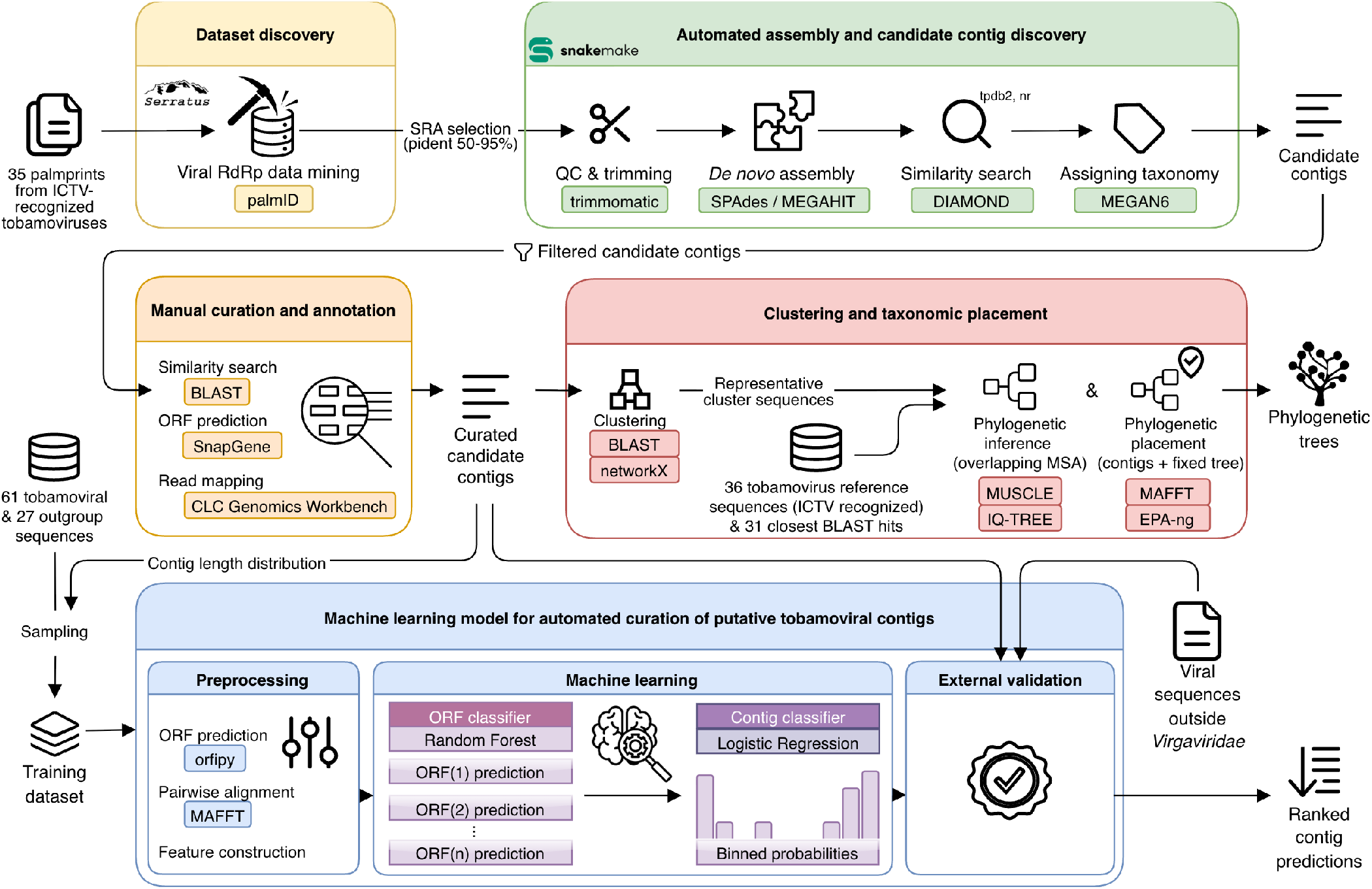
Overview of the workflow for discovery and characterization of sequences of novel tobamoviruses. The workflow is organized into five functional modules: Dataset discovery, a Snakemake-based pipeline for automated assembly and candidate contig discovery, Manual curation and annotation, Clustering and taxonomic placement, and the Machine learning model each module, colored boxes indicate the software used to perform the corresponding steps. Abbreviations: ICTV - International Committee on Taxonomy of Viruses, SRA - Sequence Read Archive, RdRp - RNA-dependent RNA polymerase, QC - quality control, tpdb2 – custommade tobamoviral protein sequence database, nr - non-redundant NCBI protein database, MSA - multiple sequence alignment, ORF - open reading frame. **Alt text:** Flowchart illustrating a five-stage viral discovery workflow. The process begins with dataset discovery via viral RNA-dependent RNA polymerase mining, followed by automated assembly and contig discovery using Snakemake. Candidate contigs then undergo parallel tracks for manual curation, clustering for phylogenetic analysis, and automated classification using a supervised machine learning model.

## Methods

### Selection of SRA datasets for re-analysis

To discover datasets containing potential novel tobamoviruses, we used the Serratus PalmID tool (Edgar et al., 2022) to obtain sequence read archive (SRA) datasets containing sequences similar to RdRp amino acid sequences from 35 of the 37 *Tobamovirus* species recognized by ICTV (obtained in March 2022), excluding tobacco latent virus (no RdRp sequence available) and maracuja mosaic virus (only partial replicase sequence available, no viral RdRp identified in input). We collated the search results, retaining only the viral hit with the highest percentage identity (pident) for each unique run ID and palm ID pair, ensuring that each candidate dataset was represented by its most informative alignment. Next, we applied a filtering step based on sequence identity, selecting hits with pident values between 50–95%. The lower bound (50%) was chosen to exclude spurious, weakly related matches, while the upper bound (95%) removed near-identical sequences corresponding to known tobamoviruses. By focusing on this range, we aimed to identify datasets that most likely contain sequences of novel tobamoviruses. We downloaded the identified datasets using fasterq-dump (v3.0.2) for downstream analyses (SRA Toolkit Development Team, 2025). Metadata for each associated SRA record were gathered using `SRAweb()` from the `pysradb` (Choudhary, 2019) and systematically recorded for subsequent comparative analysis, see Supplementary Table S4. Custom dataset selection is available online (Pajek Arambašič et al., 2026a).

### Snakemake pipeline for the automated assembly and discovery of putative tobamoviral candidate contigs

We developed a Snakemake pipeline (Mölder et al., 2021) to automate the initial processing steps, including read assembly from short-read datasets, sequence similarity search against known tobamoviral and other sequences, and filtering steps that enrich for contigs likely to represent novel tobamoviruses. Within the Snakemake pipeline, downloaded reads were first trimmed with Trimmomatic (v0.39) (Bolger et al., 2014) and assembled into contigs using SPAdes isolate (v3.15.4) (Bankevich et al., 2012) or MEGAHIT (v1.2.9) (Li et al., 2015) when SPAdes failed. Assembled contigs were filtered by length, retaining sequences between 600 and 8000 nucleotides to capture complete or partial tobamoviral genomes (typically 6300 to 6500 nucleotides) while excluding short fragments and non-targeted long sequences. Filtered contigs were first searched for similarity using DIAMOND blastx (v0.9.14) (Buchfink et al., 2021) against a local tobamoviral database (tpdb2). The tpdb2 comprised 2,177 protein sequences retrieved from the NCBI Identical Protein Groups using the query “tobamovirus” with “viruses” division filter (downloaded January 4, 2022). Contigs with hits to tpdb2 were subsequently searched against the NCBI non-redundant database (nr, built January 4, 2022, using DIAMOND blastx). Taxonomic assignment was performed using Megan6 (v6.21.7) (Beier et al., 2017) with the lowest common ancestor (LCA) algorithm. Contigs classified as cellular organisms in the second DIAMOND blastx search were removed from further analysis.

Following the Snakemake pipeline, the resulting contigs were further inspected and filtered to ensure data quality and consistency. Duplicated contigs arising from multiple SRA submissions of the same dataset were identified and removed. We inspected each SRA dataset for the number of contigs and the distribution of contig lengths. Datasets displaying anomalous characteristics, such as an unusually high number of contigs, extreme contig-length distributions, or evidence of chimeric sequences, were excluded from downstream analyses. This procedure resulted in an initial list of putative tobamoviral contigs which were further manually curated by a domain expert as described below and analyzed using a newly developed supervised machine learning model.

### Manual curation, labeling, and annotation of putative tobamoviral candidate contigs

We manually inspected the initial list of putative tobamoviral contigs generated by the Snakemake pipeline described above to create a curated list of known and novel tobamoviral sequences. We performed separate blastn (NCBI nt, April 2025, `e-value: 0.05`, `word size: 11`, `gap open: 5`, `gap extend: 2`) and blastx (NCBI nr, April 2025, `e-value: 0.05`, `word size: 5`, `gap open: 11`, `gap extend: 1`, `matrix: BLOSUM62`) searches to assess local BLAST-based sequence identities to known tobamoviruses. We classified each contig into one of the six predefined categories: tob1 – sequences of known tobamoviruses, tob2 – sequences of novel tobamoviruses, tob3 – sequences of highly divergent potentially novel tobamoviruses, where it was difficult to assign the genus based on sequence similarity of contigs representing partial genomes, oth1 – sequences related to members of other genera from *Virgaviridae* family, oth2 – sequences belonging to other viruses not related to *Virgaviridae* family, and mas – misassembled sequences. For putative misassembled contigs, we performed read mapping with CLC Genomics Workbench (v25.0.3, QIAGEN) with the following parameters: `length fraction: 0.9`, `similarity fraction: 0.9`, `match score: 1`, `mismatch cost: 1`. We visually inspected mappings to confirm that the contigs were misassembled. For tob3 contigs (highly divergent from recognized tobamoviruses), we also examined genome organization to evaluate consistency with the typical genome architecture for tobamoviruses.

We predicted open reading frames (ORFs) for contigs of putative novel tobamoviruses (categories tob2 and tob3) with SnapGene (v7.2.1) (Dotmatics, 2026) using the following parameters: `minimum ORF length: 75`, requiring a start codon ATG, except at DNA ends. For each predicted ORF, we conducted a blastp search against the non-redundant protein database (NCBI nr, July 2025, with parameters as listed above) and used the results for contig annotation. We determined the completeness, length, and presence of the start and stop codons for each predicted ORF (Supplementary Table S3). We documented any sequence abnormalities (*e*.*g*., lack of STOP codon, ambiguous START codons, interrupted sequence), possibly caused by sequencing or assembly errors, see Supplementary Table S3 column “notes”.

### Machine learning model for automated curation of putative tobamoviral contigs

We developed a supervised machine learning model aiming to automate the curation of putative tobamoviral contigs. In practice, the model learns from known tobamoviral sequences (positive examples) and closely related *Virgaviridae* sequences (negative examples) to distinguish between them. We summarized the machine learning workflow in the following paragraphs. We also included a more detailed description of each step in the Supplementary Methods. All feature extraction scripts, cross-validation workflows, and trained models are available online (Pajek Arambašič et al., 2026a).

#### Reference sequence selection and creation of the training set

We downloaded complete genome sequences of all NCBI *Virgaviridae* members (accessed January 20, 2025). We clustered them by 90% pairwise identity using MAFFT (v7.520) (Katoh and Standley, 2013) alignments and single-linkage clustering in SciPy (v1.14.1) (Virtanen et al., 2020), retaining one representative per cluster to construct a reference database of 61 tobamoviral and 27 outgroup sequences (representatives of other *Virgaviridae* members, see Supplementary Table S6). To increase data diversity under realistic conditions, we applied data augmentation by generating synthetic contigs that mimic fragmented assemblies. From each reference genome, we randomly sampled 30 contigs based on the observed length distribution of the candidate tobamoviral contigs obtained via the Snakemake pipeline and manual curation. For multipartite genomes, we selected fragments in proportion to their genomic length. We labeled all synthetic contigs by source genome as tobamoviral or outgroup.

#### Sequence alignment with reference ORFs, alignment, and sequence-based features

To construct features for contig machine-learning classification, we derived both alignment-based and sequence-based features from each predicted ORF. To predict partial and complete ORFs, we used orfipy (Singh and Wurtele, 2021) and an additional six-frame translation to ensure detection of coding regions lacking start or stop codons. We aligned each predicted ORF against reference amino acid sequences using MAFFT (v7.520) (Katoh and Standley, 2013), excluding sequences from the same parent genome. Of the four viral proteins initially considered (RdRp ORF1, RdRp ORF2, CP, and MP), we excluded MP because these proteins are possibly not of monophyletic origin across all *Virgaviridae* members (Butković et al., 2023). We computed pairwise alignment metrics, as detailed in Supplementary Table S1. We aggregated alignment statistics across reference sequences by protein (ORF1, ORF2, or CP) and reference group (tobamoviral *vs*. outgroup), using the minimum, mean, and maximum values, yielding 144 alignment-derived features per ORF. In addition, four sequence-based features were included (ORF length, contig lengths, stop codon count, and ORF prediction method). The final dataset comprised 148 features for 34,492 predicted ORFs. Model training used forward-strand ORFs, excluding RNA segments encoding only MP or triple gene block. We have provided the exact inclusion criteria in Supplementary Table S6 (column “training”). Testing was performed using ORFs from all genomic segments, including reverse-strand ORFs.

#### Model selection for ORF classifier

We trained and compared several supervised learning algorithms to identify the most effective model for distinguishing tobamoviral and non-tobamoviral ORFs. These included Logistic Regression, Support Vector Machines, Decision Trees, Naive Bayes, K-nearest Neighbors, and Random Forest, implemented in scikit-learn (Pedregosa et al., 2011). We performed model selection using a repeated stratified cross-validation framework (5 iterations × 5 folds). From this process, we selected Random Forest with 200 estimators and a maximum depth of 50 to predict whether a given ORF is a tobamovirus or an outgroup. For feature interpretation, we scored and ranked feature importance using the Random Forest ORF classifier trained on the reference training dataset, quantified by scikit-learn’s mean decrease in impurity (MDI) importance score. We additionally performed a feature-enrichment analysis to test whether predefined feature classes were overrepresented among the top-ranked features (the top 40 of 148) compared to their background frequency, using one-sided hypergeometric tests with Bonferroni correction for multiple comparisons. For details, see Supplementary Methods S1.

#### Contig classifier, aggregation strategies of ORF-level predictions using stacked generalization

Each contig may include a varying number of individual ORF predictions. Because biological interpretation requires a single prediction per contig, ORF-level predictions were aggregated into a contig-level classification using stacked generalization, as introduced in Wolpert (1992). We evaluated two aggregation strategies: (i) **Extreme classifier**, which assigns the contig label based on the ORF with the probability farthest from the 0.5 classification threshold, and (ii) **Binned classifier**, using a logistic regression meta-classifier applied to a vector of binned ORF probabilities (10 bins). Additional binning schemes were also tested, see Supplementary Methods and Supplementary Figure S3. We compared both aggregation strategies using cross-validation and calibration analyses.

#### Evaluation of the contig classifier

We assessed contig-level performance of both aggregation strategies using 30 independent 5-fold stratified cross-validation iterations (150 total evaluations). Within each fold, ORF-level predictions followed a leave-one-virus-out strategy to prevent sequences from the same viral species from appearing in both training and test sets. We evaluated performance using accuracy, F1 score, precision, recall, and AUC with 95% confidence intervals estimated across iterations. We assessed probability calibration for both aggregation strategies using 10-bin reliability curves computed with `sklearn.calibration.calibration_curve`, pooling predictions across folds and summarizing curves relative to the ideal diagonal line. In addition, we performed a threshold analysis for the selected Binned classifier to evaluate fixed versus tuned decision thresholds. Further details and results are provided in Supplementary Figure S4.

We further evaluated the model robustness and its ability to reject non-*Virgaviridae* sequences using two external tests designed to reflect realistic conditions. First, to assess discrimination against more divergent viruses, we analyzed 1,398 NCBI viral representatives (accessed January 16, 2025) deduplicated at the genus level by running relevant parts of the Snakemake pipeline (specifically, DIAMOND against the tpdb2 database). The 41 sequences that passed the prefilter were evaluated by the model to estimate the false positive rate. Second, we applied the model to candidate tobamoviral contigs obtained by data mining in this study and compared the predictions with those from expert manual curation.

### Clustering and phylogenetic analyses for contigs of putative novel tobamoviruses

We focused on further characterizing the putative novel tobamoviral sequences identified in this study through data mining. We clustered those sequences (categories tob2 and tob3, from the manual curation process described above) using a custom pipeline (Pajek Arambašič et al., 2026a) (Figure 1, “clustering”). First, an all-vs-all blastn was performed using the BLAST+ command line utility (v2.16.0) with the following parameters: `e-value: 10`, `word size: 11`, `gap open: 5`, `gap extend: 2`, `match reward: 2`, `mismatch penalty: -3`. The blast output table was deduplicated and filtered to remove self-hits, retaining only hits with at least 90% identity spanning at least 50 nucleotides. From the remaining hits, a network graph was built with NetworkX (v3.2) (Hagberg et al., 2007), with nodes representing contigs and edges representing sequence similarity. Contigs within each connected component were assigned to the same cluster using the NetworkX function `nx.connected components`.

First, we built separate phylogenetic trees for each of the four tobamoviral ORFs. For each ORF, cluster members were aligned with MUSCLE in MEGA11 (v11.0.13) (Tamura et al., 2021), and one representative per cluster was selected to be included in the analysis (the sequence spanning the longest region aligned across most contigs). Cluster representatives were queried against the NCBI nr database using blastx (`e-value: 0.05`, `word size: 5`, `gap open: 11`, `gap extend: 1`, `matrix: BLOSUM62`) and the top 10 unique hits (E-value) per representative were aligned together (as described above) with cluster representatives, reference tobamoviral sequences from the ICTV VMR table (MSL40, v1.20250307), and outgroup sequences. Oat golden stripe virus (*Furovirus*) served as the outgroup for the RdRp ORF1, RdRp ORF2, and MP trees, and pepper ringspot virus (*Tobravirus*) for CP. We discarded sequences that did not align to the region shared by most representatives, manually trimmed the ends to the longest aligned segment, and applied further automated trimming with Trimal (v1.13, method `automated1`) (Capella-Gutiérrez et al., 2009) (files available as Supplementary Files S1-S4). Maximum-likelihood trees were then inferred from the trimmed alignments in IQ-TREE (v3.0.1) (Nguyen et al., 2015), using automatic substitution-model selection and bootstrap support from the bootstrap splits method (default parameters).

To obtain a single phylogeny for all cluster representatives, we also performed phylogenetic placement with EPA-ng (v0.3.8) (Barbera et al., 2019) on a precomputed reference phylogenetic tree built from the same reference dataset used for ORF-based phylogenetic trees. We aligned reference sequences with MAFFT (Katoh and Standley, 2013) and used the untrimmed alignment to infer a fixed maximum-likelihood reference tree with IQ-TREE (v3.0.1) (Nguyen et al., 2015) (`-m MFP`, `-bb 1000`). We aligned each query contig individually to the fixed reference multiple-sequence alignment using MAFFT (`--addfragments --keeplength`), and retained only the aligned fragment to preserve the reference alignment coordinates. We then combined query-aligned fragments and placed them jointly onto the fixed reference tree using the corresponding substitution model in a single EPA-ng run. We stored placement outputs in .jplace files and subsequently analyzed them with gappa (v0.9.0) (Czech et al., 2020) for downstream summarization and interpretation of phylogenetic placement. We used iTOL (v7.2.2) (Letunic and Bork, 2021) for visualization and annotation of phylogenetic trees.

To estimate the minimum number of discovered novel tobamoviruses, we identified, for each clade of novel viruses on the EPA-ng-derived tree, the largest set of contigs sharing at least a 50-nt mutually overlapping region in the reference alignment. We used the size of this maximal-overlap set as a lower-bound estimate of the number of distinct viruses.

## Results

We initially identified 6,870 candidate sequencing runs (SRRs) using the Serratus search. To focus on the discovery of novel tobamoviruses, we filtered datasets based on alignment identity thresholds, retaining only SRRs with palmID hits and a pairwise identity in the range of 50%–95%, yielding 253 SRRs.

### The developed Snakemake pipeline narrows the set of candidate tobamoviral contigs

All selected SRRs were processed using the Snakemake pipeline, yielding 2,567 assembled contigs from 145 SRRs. We excluded 166 contigs previously annotated as derived from cellular organisms and 18 contigs (6 SRRs) identified as duplicates due to multiple SRA submissions of identical datasets deposited under different accession numbers. Next, we assessed assembly quality and dataset-specific features (contig number and size distributions, and the presence of chimeric sequences) across selected SRRs. After inspecting the assembly features, we decided to exclude contigs from the dataset SRR6846476. This dataset comprised 1,873 contigs and displayed a markedly different composition compared to the others. Manual examination revealed numerous chimeric sequences and very short contigs (∼600 nt) near the lower length threshold applied in the Snakemake pipeline. Given these anomalies and the potential to introduce artifacts, we excluded this dataset from downstream analyses. After filtering, 510 contigs from 131 SRRs remained for manual curation and labeling.

### Manual curation confirms 45% of identified contigs are tobamoviral

Each contig was assigned to one of six predefined categories, as described in the Methods section. Out of 510 contigs, we classified 228 contigs (45%) as tobamoviral (categories tob1, tob2, and tob3) and assigned the rest to the remaining categories (oth1, oth2, mas). A summary of the manual curation, including notable observations on tobamovirus-like contigs and other viral groups, is provided in Table 1.

**Table 1.** Summary of manual curation of 510 discovered contigs from this study with the corresponding number of source datasets (SRRs).

| Manual curation category | Abbreviation | Number of contigs | Number of SRR datasets |
| --- | --- | --- | --- |
| Sequences of known tobamoviruses | tob1 | 83 | 13 |
| Sequences of novel tobamoviruses | tob2 | 109 | 15 |
| Sequences of highly divergent potentially novel tobamoviruses | tob3 | 36 | 8 |
| Sequences related to other genera from *Virgaviridae* family | oth1 | 109 | 65 |
| Sequences belonging to other viruses not related to *Virgaviridae* family | oth2 | 151 | 77 |
| Misassembled sequences | mas | 22 | 10 |

### Machine learning approach aids in data curation step

To automate contig curation, we developed a supervised machine-learning approach to predict whether a contig is tobamoviral. The model first scores each ORF on a contig. It then combines the ORF scores into a single classification for the entire contig.

For the ORF-level classification, we tested six classifiers. Random Forest performed best in 22 out of 25 cross-validation folds (88%) with a mean accuracy of 0.767 ± 0.021, outperforming the second-best SVM algorithm, which was selected in only 3 folds (12%) with a mean accuracy of 0.752 ± 0.014.

The final Random Forest ORF classifier did not rely on a single dominant feature. The most influential features were primarily alignment-derived metrics (identity, gap, and alignment-length descriptors) and the number of stop codons, which provides information about the strand and the correct open reading frame. Importance was balanced between tobamovirus- and outgroup-derived features and similarly distributed across ORF1, ORF2, and CP feature groups (Supplementary Table S2). Features based on sequence identity and alignment gaps were overrepresented among the top 40 most important features (Supplementary Figure S2).

Although the ORF-level classifier performed moderately, combining the predictions from all ORFs on a contig substantially improved performance. We tested two methods to combine ORF scores. The Binned classifier consistently outperformed the Extreme classifier (Figure 2a,b). Binned contig classifier achieved an average accuracy of 0.9735 (95% CI: 0.9727–0.9743), F1 score of 0.9806 (95% CI: 0.9801–0.9812), high precision (0.9925), recall (0.9690), and AUC of 0.9971 (95% CI: 0.9969–0.9972). By contrast, the Extreme classifier reached an accuracy of 0.9231 (95% CI: 0.9217–0.9246), an F1 score of 0.9475 (95% CI: 0.9465–0.9484), and a lower precision (0.9003), despite a recall close to 1.0. The Binned classifier’s predicted probabilities closely matched the observed outcome frequencies, indicating near-perfect calibration and no need for threshold tuning (Supplementary Figure S4, Supplementary Methods S2).

**Fig. 2:**
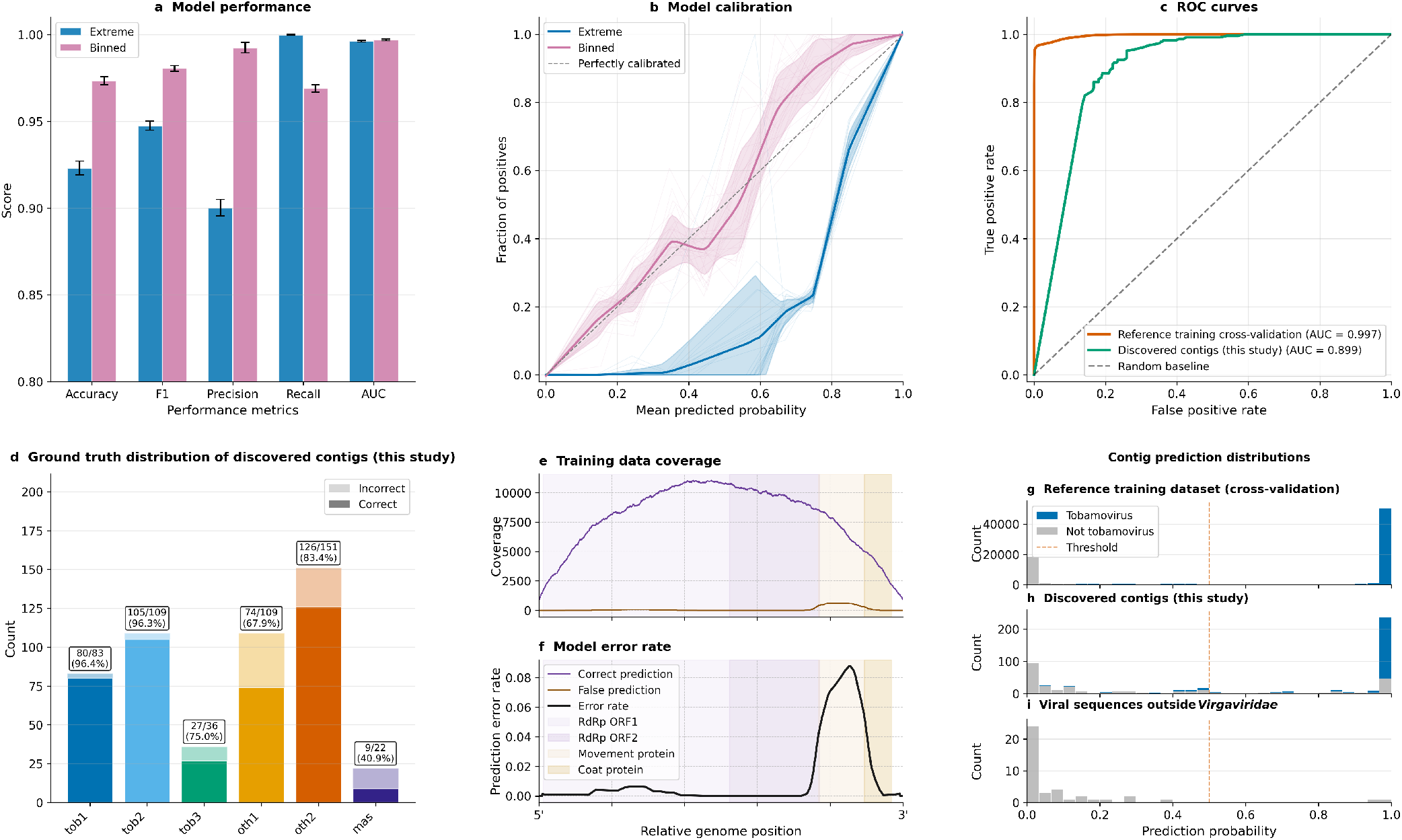
Comprehensive cross-validation and performance evaluation for the tobamovirus classification model. (a-d) Model evaluation summary showing method-wise performance metrics comparison across accuracy, F1-score, precision, recall, and ROC-AUC (a), probability calibration against the ideal diagonal (b), ROC curves for model performance on cross-validation dataset (30 fold average) and on 510 contigs discovered in this study (single fold) with AUC values (c), and ground-truth category composition with correctly classified contigs (solid bars) and misclassified contigs (transparent overlays), including per-category sample sizes and accuracies. X-axis category abbreviations are defined in Table 1 (d). (e,f) Training data coverage (e) and model error rate (f) with relative genome position, RdRp - RNA-dependent RNA polymerase, ORF - open reading frame. (g-i) Contig prediction distributions with Tobamovirus sequences in blue for three datasets: Reference training dataset (g), discovered contigs from this study (h), and Viral sequences outside Virgaviridae (i), with the decision threshold indicated by a dashed vertical line at (p = 0.5) separating outgroup-like (p < 0.5) and tobamovirus-like (p ≥ 0.5) predictions. **Alt text (Panel a):** Grouped bar chart comparing extreme and binned model performance across five metrics. The binned configuration shows higher scores for accuracy, F one, and precision, while the extreme configuration has higher recall. All scores are above zero point eight. **Alt text (Panel b):** Calibration plot showing the relationship between predicted probability and the fraction of positives. The binned model variant follows the perfectly calibrated diagonal more closely than the extreme model variant. **Alt text (Panel c):** Receiver operating characteristic curves for two datasets. The reference training cross-validation achieves an area under the curve of zero point nine nine seven, while the discovered contigs achieve zero point eight nine nine. **Alt text (Panel d):** Stacked bar chart showing the ground truth distribution of discovered contigs across six categories, with each bar split into correctly and incorrectly classified contigs. Labels indicate the percentage of correct predictions, highest for known tobamoviral sequences at ninety-six point four percent, followed closely by novel tobamoviruses at ninety-six point three percent, and lowest for misassembled contigs at forty point nine percent. **Alt text (Panel e):** Line graph showing sequence coverage depth across the relative genome position. Coverage values rise from approximately two thousand at the ends to over ten thousand in the center of the genome. **Alt text (Panel f):** Line graph showing the model error rate relative to genome position. Shaded regions identify genes for RNA-dependent RNA polymerase, movement protein, and coat protein, showing a higher error rate in the movement protein region. **Alt text (Panel g):** Histogram showing prediction probability distributions for the reference training dataset. A vertical dashed line at zero point five marks the threshold, with most sequences clustered at zero point zero and one point zero. **Alt text (Panel h):** Histogram showing the prediction probability distribution for discovered contigs. The majority of sequences in this study are correctly predicted as viral with probabilities near one point zero. **Alt text (Panel i):** Histogram showing the prediction probability distribution for viral sequences outside the target family. Most of these sequences correctly fall below the classification threshold of zero point five.

To evaluate the model’s performance on highly divergent viral sequences (outside the *Virgaviridae* sequence space used for model training), we further applied the model to 41 selected non-redundant viral sequences, see Methods. Of those 41, 39 were assigned to the outgroup, and 2 to the tobamovirus category.

The final task of the model was to evaluate its predictive ability on the curated set of 510 contigs derived from the Snakemake pipeline data mining output from this study. We compared model predictions with manually curated annotations from domain experts, enabling a realistic assessment of model performance. This evaluation considered both overall accuracy and per-category agreement, focusing on tobamovirus-like contigs, other viral groups, and misassembled sequences.

Across the entire dataset, the model achieved strong performance (AUC-ROC 0.899), indicating good discriminative ability between tobamovirus-like and non-tobamoviral sequences. Out of 510 contigs, 212 were classified as true positives, 73 as false positives, 209 as true negatives, and 16 as false negatives. These values correspond to a precision of 74.4%, a recall of 93.0%, and an F1-score of 82.7%.

When broken down by category, model accuracy was highest for tobamovirus-like groups tob1 (96.4%) and tob2 (96.3%), while somewhat reduced performance was observed in tob3 (75.0%), likely reflecting higher sequence divergence for this group. Among other viral categories, oth1 contigs showed moderate accuracy (67.9%), whereas oth2 contigs were classified with higher reliability (83.4%). In contrast, the misassembled sequence category (mas) posed the greatest challenge, with the model correctly classifying only 40.9% of contigs.

### Novel tobamoviral contigs are associated with diverse dataset types

The highest number of datasets in which we detected sequence similarities with tobamoviruses using the Snakemake pipeline was associated with plant-related material (40 in total), followed by 36 from animal-related and 29 from human-related samples (Figure 3a). However, further analysis revealed that most sequences in these datasets corresponded to other RNA viruses (oth1, oth2) rather than tobamoviruses. After manual review, we identified sequences classified as known and novel tobamoviruses in 23 SRRs, of which 10 were from freshwater environments, all deposited as metatranscriptomic datasets of samples from the Amazon River (PRJNA237345). These SRRs yielded the large majority of detected clusters of putative novel tobamoviruses (Figure 3b), for example, cluster 4 (14 contigs) and cluster 10 (4 contigs), for which the phylogenetic placement is described in detail below.

**Fig. 3:**
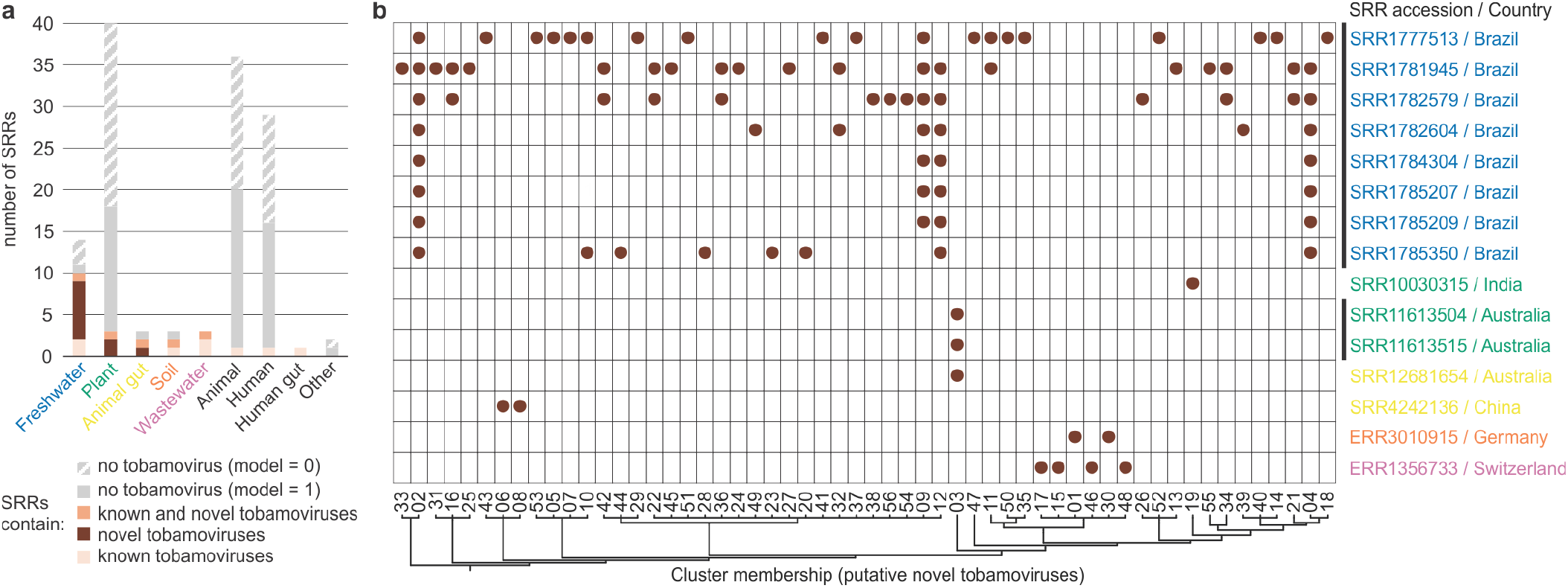
SRR datasets containing discovered contigs (this study) with detected tobamoviral sequence similarities and their associated sample types. (a) Number of SRRs associated with organism (e.g., animal, human, plant) or environment (e.g., freshwater, wastewater, soil) sample types, split by presence/absence of known or novel tobamoviral contigs (as defined by manual curation). Category “no tobamovirus” is broken into two parts, separating SRRs in which the machine learning model did not recognize any tobamoviral contigs (model=0) or such that the model falsely predicted some contigs as tobamoviral (model=1). (b) Presence of putative novel tobamoviral clusters (columns) across individual SRRs arranged by sample type (lines). Dot denotes detection of a cluster in the corresponding SRR, colors of the SRR accessions represent the sample type as defined in (a), black lines next to the SRR accessions connect samples from the same study. Viral clusters are ordered according to their phylogenetic placement relationships, as denoted by the dendrogram below the panel. **Alt text (Panel a):** Bar chart displaying the number of sequence read archive runs across nine sample categories. Freshwater and plant-associated datasets show the highest counts of novel tobamoviruses. Shaded bar segments differentiate sequences by curation category, including known viruses, novel viruses, and those with no detected tobamoviruses. **Alt text (Panel b):** Matrix plot showing the distribution of novel tobamovirus clusters across various sequence read archive datasets. Accessions are labeled by country of origin, including Brazil, India, Australia, China, Germany, and Switzerland. Dots indicate the presence of specific viral clusters, which are organized horizontally according to their phylogenetic relationships, as shown in the dendrogram. The majority of dots are from Brazilian freshwater datasets.

Although we analyzed only a small number of wastewater-associated SRRs, they contained a relatively high proportion of tobamoviral contigs (43 out of 228 in total), the majority of which corresponded to known tobamoviruses (39 contigs). Putative novel tobamoviral sequences identified in the wastewater metagenome from Switzerland were part of four wastewater-associated clusters (15, 17, 46, and 48). Furthermore, we detected sequences of potentially novel tobamoviruses from plant-associated transcriptomes (*Nicotiana tabacum*, Saccharum hybrid) (clusters 3 and 19), animal gut datasets (clusters 3, 6, 8), and soil microbiomes (clusters 1 and 30).

Most of the clusters representing putative novel tobamoviruses contained contigs from the same studies and, therefore, from the same type of samples. Cluster 3 contained contigs from the same country but different sample types, including animal gut (Australian tropical cattle rumen microbiome) and plants (sugarcane).

### Phylogenetic analyses reveal clustering of novel tobamoviral sequences within subgroups across the genus

We constructed maximum likelihood phylogenetic trees for the four tobamoviral ORFs to determine how representatives of the clusters containing putative novel tobamoviruses (categories tob2 and tob3) relate to known tobamoviruses (Figure 4b,c,d,e, Supplementary Figures S6, S7, S8, S9). Additionally, we performed phylogenetic placement of the representatives into a single reference tree to enable comparison of sequences covering different parts of the genome (Figure 4a). Phylogenetic analysis showed that cluster representatives grouped with different clades within the genus *Tobamovirus* (Figure 4, Supplementary Figures S6, S7, S8, S9). Overall, the results from the phylogenetic trees based on the four ORFs were consistent with the results of the phylogenetic placement, with occasional discrepancies for individual cluster representatives. Some notable examples of grouping within the *Tobamovirus* genus, based on the phylogenetic placement tree, are highlighted below.

**Fig. 4:**
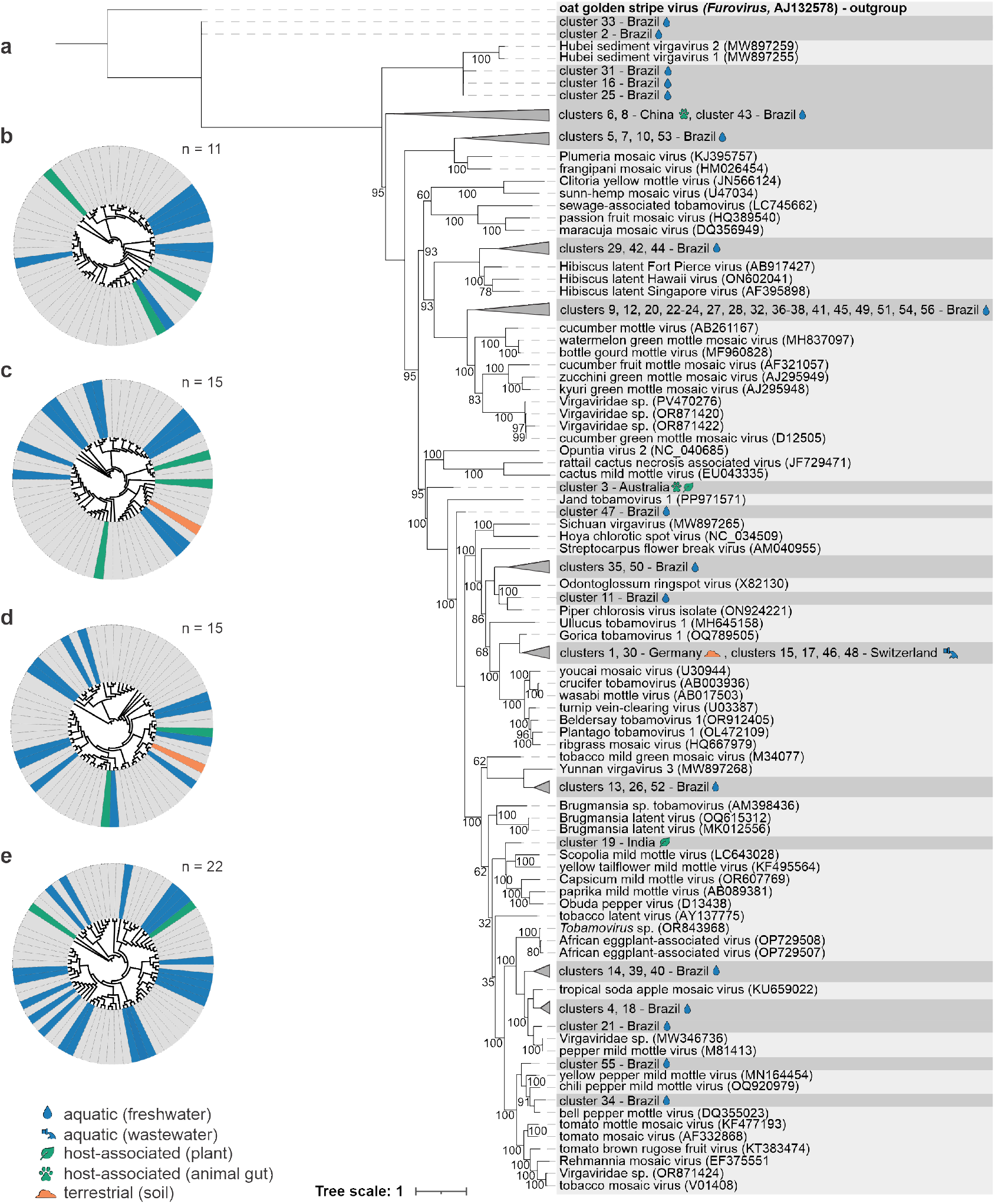
Phylogenetic analysis of novel tobamoviral sequences. (a) Phylogenetic placement of cluster representatives onto a maximum likelihood reference tree. The reference tree was constructed from complete genomic sequences of the closest blastx hits of the cluster representatives and the ICTV VMR tobamoviral reference sequences using the GTR+F+R6 substitution model. The tree was rooted with an outgroup (oat golden stripe virus, genus Furovirus), shown in bold. Support values derived from the ultrafast bootstrap analysis are displayed on the branches and refer only to the reference tree topology. Cluster representatives of novel tobamoviruses were subsequently placed onto the reference tree with the EPA-NG phylogenetic placement tool. For each cluster representative, the country of origin of the dataset used to assemble the contigs is indicated, along with an icon representing the source material type, as defined in the legend below. GenBank accessions for cluster representatives of novel tobamoviruses are listed in Supplementary Table S5, column “genbank accession number”. (b-e). Schematic representation of the maximum-likelihood phylogenetic trees inferred from the nucleotide sequences of RdRp ORF1 (b), RdRp ORF2 (c), CP (d), and MP (e). The colors correspond to the source material from which the datasets used for assembling each cluster’s contigs originate, as shown in the legend below. The number of novel tobamoviral contigs included in each tree is denoted on the upper right side. **Alt text (Panel a):** Phylogenetic tree constructed with phylogenetic placement of novel tobamoviruses on the maximum likelihood reference tree, inferred from reference tobamoviral sequences and closest blastx hits of cluster representatives. Most cluster representatives are grouped within the main tobamovirus clade but distributed across several distinct subgroups. **Alt text (Panels b-e):** Schematic representation of maximum likelihood phylogenetic trees inferred from nucleotide sequences of the four tobamoviral open reading frames. The branches are colored by source material (freshwater, wastewater, plant, animal gut, soil). The number of novel tobamovirus contigs is indicated for each tree, with the tree based on open reading frame four containing the highest number of contigs.

Using the overlap-based counting approach described in the Methods section, we estimated that novel tobamoviral sequences correspond to a minimum of 32 distinct novel tobamoviruses across the identified EPA-ng clades. This value represents a lower-bound estimate under the assumption of non-recombinant genome structure.

Cluster 4 was among the largest analyzed clusters, containing 14 contigs detected in 7 datasets from the Amazon River study (Supplementary Table S3). It formed a distinct clade, together with tropical soda mosaic virus (TSAMV) and pepper mild mottle virus (PMMoV), in both the phylogenetic placement tree (Figure 4) and individual phylogenetic trees inferred from ORFs 1–4 (Supplementary Figures S6, S7, S8, S9). Several other clusters with contigs derived from the same study were placed into the same group in the phylogenetic placement tree, including clusters 14, 18, 21, 39 and 40 (Figure 4a).

Cluster 10 consistently represented a sister clade to the clade containing Plumeria mosaic virus and frangipani mosaic virus across all analyzed trees (Figure 4a, Supplementary Figures S7, S8, S9). Several other representatives, including those from clusters 5, 7 and 53, were also placed in the same clade in both the phylogenetic placement tree (Figure 4a) and individual trees based on ORFs 1, 2 and 4 (Supplementary Figures S6, S7, S9).

Representatives from clusters 1, 15, 17, 30, 46, and 48, together with Gorica tobamovirus 1, formed a sister clade in the phylogenetic placement tree to a group of known tobamoviruses, including ribgrass mosaic virus, turnip vein-clearing virus, wasabi mottle virus, youcai mosaic virus, Plantago tobamovirus 1, and crucifer tobamovirus (Figure 4a). Similarly, a large set of cluster representatives (N = 18) was grouped with viruses that can infect members of the *Cucurbitaceae* family, as well as three novel tobamoviruses designated as *Virgaviridae* sp. (Figures 3, 4a).

With 35 contigs, cluster 2 is the largest in our dataset (from 8 datasets derived from the Amazon River study). We assigned the contigs from this cluster to the category tob3 due to low sequence identity with known tobamoviruses. The representatives of cluster 2 and 33 represent a basal lineage on the inferred phylogenetic placement of the tobamovirus tree, as well as other tob3 clusters (16, 31, 33) that group with Hubei sediment virgavirus 1 and 2 (Figure 4). The basal position of these viral sequences is not supported in separate ORF trees (see Supplementary Figures S6, S8, S9), where the representatives of clusters 2, 31, and 33 fall within the known tobamoviral clades, often with long branches indicating substantial genetic divergence.

## Discussion

### Discovery of a diverse set of previously unknown tobamoviruses across environmental sample types

Our results reinforce the expectation that tobamoviruses can be detected across a wide range of environmental samples, consistent with their observed environmental stability and persistence (Mehle et al., 2018). By mining publicly available sequencing datasets originally generated to address diverse biological questions, we identified tobamoviral sequences in freshwater, wastewater, plant transcriptomes, animal gut microbiomes, and soil samples.

Freshwater environments, particularly the Amazon River metatranscriptome, were the predominant source of novel tobamoviral clusters, which, in the majority of cases, grouped within established tobamoviral subclades. Clusters originated from various locations along the Amazon River, including one pristine site (Topajos: SRR1777513) and three sites adjacent to urban areas (Obidos: SRR1781945, SRR1782579, SRR1782604, Macapa North Channel: SRR1785209, SRR1785350, Macapa South Channel: SRR1784304, SRR1785207). The dataset from the pristine site contained the highest number (16) of unique novel tobamoviral clusters (*i*.*e*., those appearing only in this dataset), which might reflect high plant diversity in the surrounding environment.

Phylogenetic analyses revealed that the putative novel tobamoviral sequences identified in this study are distributed across multiple major clades within the genus. The phylogenetic placement approach employed in this study enabled the integration of partial contigs spanning different genomic regions into a unified phylogenetic framework, facilitating the estimation of the minimal number of distinct novel viruses and the identification of relationships among clusters. Using conservative overlap-based estimates, we inferred a minimum of 32 distinct novel tobamoviruses from 15 datasets across 7 studies (Table 1, Figure 3), substantially expanding the known diversity of the genus (37 ICTV-recognized species, as of July 2026). Several clusters formed well-supported sister clades to recognized tobamovirus species groups, including those related to PMMoV and cucurbit-associated viruses, capturing diversity closely related to agriculturally relevant lineages.

Notable novel tobamovirus findings from Amazon River datasets include 18 clusters that group with known cucurbit-associated tobamoviruses and three novel species (*Virgaviridae* sp.) previously identified in metagenomic datasets from bat intestine (China) and tunicate (Australia) (Petrone et al., 2025). Their consistent placement within a clade infecting agriculturally important crops suggests that these clusters may represent novel viral species with potential agricultural relevance.

The frequent detection of tobamoviral sequences in freshwater samples in this study (see Figure 3), despite their geographic restriction to the Amazon River, highlights these environments as important sample types for the discovery of stable viruses. These findings are consistent with previous reports demonstrating the persistence and high abundance of tobamoviruses in aquatic matrices and further support the idea that freshwater systems may integrate viral inputs from diverse terrestrial and plant-associated sources (Zhang et al., 2022; D’Aoust et al., 2026; Maksimović et al., 2024).

Wastewater samples were also a notable source of tobamoviral diversity. Although only a small number of wastewater-associated SRRs were analyzed, they contained a substantial number of tobamoviral contigs, including several cluster representatives most closely related to *Gorica tobamovirus 1*, which was previously identified in various irrigation water sources in Slovenia. Viruses from this clade were previously frequently detected in environmental water samples, shown to infect several wild plant species (such as ribgrass), but, in contrast with some other known tobamoviral clades, were not linked with outbreaks in solanaceous crops (Maksimović et al., 2024). All contigs from these clusters originated either from wastewater (15, 17, 46, 48) or silage digestate (1, 30), both collected in Europe (Germany, Switzerland), in contrast to most other clusters, which were associated with freshwater samples from Brazil. The presence of tobamoviruses in wastewater is consistent with the high stability of tobamoviral particles, which can pass through human gastrointestinal tracts as viable virions (Colson et al., 2010). Pepper mild mottle virus (PMMoV) exemplifies this property and is widely used as an indicator of human fecal load (Rosario et al., 2009; Kitajima et al., 2018). Thus, wastewater effectively integrates viral signals from human consumption, agricultural produce, and environmental inputs, making it a valuable matrix for detecting both known and emerging plant viruses (Bačnik et al., 2020; Tisza et al., 2023).

Most clusters of putative novel tobamoviruses were associated with a single study or sample type, except cluster 3, which included contigs recovered from both animal gut (bovine) and plant-associated (sugarcane) datasets from the same country (Australia), where sugarcane and sugarcane-derived products are sometimes used as cattle feed (Northern Australian Dairy Hub, 2018), suggesting that the detection of the virus in bovine gut could be of dietary origin.

### Highly divergent tobamoviral sequences

All contigs that we flagged as highly divergent tobamoviruses during manual curation (tob3, clusters 2, 16, 25, 31, 33) were also detected in the Amazon River datasets, mainly at sites influenced by urban activity (Obidos, Macapa South and North channels). They had the highest identity with two unclassified members of the *Virgaviridae* family discovered in lake sediment samples from China (Hubei sediment virgavirus 1, 2) (Chen et al., 2022). While these two viruses were not classified as tobamoviruses in the original study, we investigated their genome structure together with that of several longer contigs recovered in this study and observed that these viruses possess a canonical genome organization characteristic only for tobamoviruses within the *Virgaviridae* family (RdRp, followed by MP, followed by CP). Based on this genome architecture, we tentatively classified them as tobamoviruses in the present study. They occupied long branches or basal positions in phylogenetic placement analysis. These sequences likely represent divergent tobamoviruses at the boundary of current known diversity within the genus. Their phylogenetic positions were not always congruent on individual ORF trees, possibly reflecting the limited phylogenetic signal available in short or fragmented sequences. These inconsistencies in phylogenetic placement highlight a fundamental constraint of environmental virus discovery: incomplete genomes can obscure evolutionary relationships and complicate taxonomic assignment, particularly for highly divergent sequences. As additional sequencing data become available, especially datasets enabling recovery of near-complete genomes, these phylogenetic relationships can be revisited and refined. As all members of this group have so far been detected exclusively in environmental samples, further investigation will be required to determine their host range.

### Other viral discoveries outside *Tobamovirus* genus

Among the 510 discovered contigs in this study, we also identified sequences showing identity to viruses previously described as virga-like viruses (*e*.*g*., Hubei virga-like virus 11 (Shi et al., 2016; Mahar et al., 2020)) or tobamo-like viruses (*e*.*g*., plant-associated tobamo-like virus 1 (Rivarez et al., 2023)), and have classified them in the oth1 category after manual curation (see Supplementary Table S3). Some of these viral sequences lack genes encoding movement proteins, and their potential association with plant hosts therefore remains uncertain. Nevertheless, they encode proteins that are apparently homologous to proteins of members of the *Virgaviridae* family, suggesting a close evolutionary relationship with plant-associated viruses from *Virgaviridae*. Related virga-like and tobamo-like viruses have previously been detected in a broad range of plant-associated (tomato (Rivarez et al., 2023), *Rubus idaeus* (Higuita et al., 2024)), and invertebrate-associated (crab (Shi et al., 2016), nematodes (Huang et al., 2025), insects (Mahar et al., 2020)) datasets, which is in line with sample types of sequences detected in this study (mite species *Panonychus citri* and *Philonome clemensella*, moth *Chilo suppressalis* but also plants *Oryza sativa, Tripogonella loliiformis*). More targeted studies focusing on these viruses will be required to elucidate their hosts and ecological roles.

Additionally, we identified sequences related to previously undescribed members of the family *Virgaviridae* in two independent public datasets. One sequence (NODE 914 length 5302 cov 97.892632 SRR8455906) was detected in several datasets of cotton transcriptomes from China (SRP180246), and another (NODE 71 length 6076 cov 12.863977 SRR7976667) in datasets from groundwater sediments of an aquifer near the Colorado River in the USA (SRP164393). In both cases, the detected sequences showed only low similarity to known *Virgaviridae* members from different genera. These results suggest that the identified highly divergent viruses may represent new genera within the family *Virgaviridae*.

### Targeted data mining enables the scalable discovery of stable RNA viruses

This study demonstrates that targeted mining of public sequencing repositories can effectively uncover hidden diversity of environmentally stable RNA viruses such as tobamoviruses. By combining a highly sensitive similarity-based screen (Serratus PalmID) with a tailored assembly and filtering workflow, we reduced thousands of candidate datasets to a manageable subset enriched for tobamoviral sequences. The resulting Snakemake pipeline is modular and adaptable, as candidate selection, assembly, filtering, and classification are implemented as discrete steps that can be readily modified for different virus families, sequencing technologies, or discovery objectives.

Tobamovirus sequences were confirmed in only a subset of datasets initially flagged by conserved domain searches (23 of 253), underscoring that detection of conserved RdRp motifs alone is insufficient to predict successful recovery of longer viral genomic sequences in downstream analyses. Nevertheless, such signals serve as an effective first-pass filter, enabling rapid triage of large public datasets and focusing computationally intensive analyses on more biologically meaningful candidates. This strategy leverages the growing volume of publicly available sequencing data and enables virus discovery at a scale impractical with *de novo* sampling alone.

However, the effectiveness of this approach strongly depends on viral biology. Tobamoviruses are particularly well suited to large-scale data mining due to their exceptional environmental stability and frequent high abundance in environmental samples. In contrast, less stable viruses are likely to be systematically underrepresented in public repositories.

### Machine learning–assisted curation: strengths, limits, and scope

In this study, we also developed a supervised machine learning framework aiming to reduce the manual effort required to curate large numbers of candidate contigs. The final contig classifier achieved high average accuracy (0.974) and recall (0.969) during cross-validation on the reference dataset. Additional diagnostic checks revealed that most of the contig mislabeling was for contigs containing MP ORF (Figure 2e,f), which was not included in the training of the model due to the lack of homologous representation across all *Virgaviridae* members. This pattern suggests we could improve the model by expanding the set of ORFs used during training.

Next, the model performed somewhat inferiorly but still robustly (AUC 0.899, F1 0.827, precision 0.744, recall 0.930) on discovered contigs in this study. The performance was strongest for contigs corresponding to known and moderately divergent novel tobamoviruses (categories tob1 and tob2), indicating that the model effectively captured conserved features of tobamoviral ORFs. Performance decreased for highly divergent (tob3) and misassembled (mas) contigs, reflecting both biological and technical limitations. Highly divergent sequences share fewer detectable alignment features with reference viruses, while misassemblies introduce artifacts that violate assumptions underlying both alignment-based and sequence-based features. These cases delineate the practical boundaries of models and highlight where expert review remains essential. These limitations reinforce the importance of pairing ML-based prioritization with upstream quality control and downstream expert review, particularly when working with heterogeneous metagenomic assemblies. Moreover, because we trained the model primarily on *Virgaviridae* sequences, its decision boundary is optimized for discrimination within that family rather than for universal viral classification. Nevertheless, we tested model performance on viral sequences outside *Virgaviridae* (Figure 2i) and observed that most of such sequences (39 out of 41, see Results) were not classified as tobamoviral, which provides confidence about the broader usability of the model.

Importantly, we optimized the model for high sensitivity in the tobamovirus category, an appropriate trade-off for large-scale discovery workflows where downstream analyses are costly. Such a bias toward sensitivity is well suited to plant pathogen discovery, where minimizing false negatives is critical, and further refinement of the reference dataset (particularly expansion of the negative class) is expected to improve specificity without compromising detection performance.

Taken together, key benefits of the machine learning approach include: accelerated screening of contigs that lack clear domain hits, consistent prioritization of a large number of candidates, and the ability to capture important alignment features besides percent identity, as shown in Supplementary Table S2 and Supplementary Figure S2. It essentially extends the work of domain experts, who might scrutinize each ORF manually, into an automated module, with the flexibility to capture informative taxonomy-related features beyond single-domain searches by including evidence from multiple ORFs. We note that new taxonomic classifiers have recently been developed for short viral contigs. For example, Zheng et al. (2025) reports that VITAP can reliably classify viral sequences as short as 1kb to genus level. Such tools could complement our approach in the future, either as alternative classifiers or as additional validation layers.

### Workflow reuse, scalability, and practical scope

The workflow developed in this study is available for reuse and exhibits different degrees of flexibility across its components. The initial Snakemake-based processing framework is largely agnostic to the downstream classification task and could be trivially reused for other virus families or discovery efforts. On the other hand, the machine-learning component relies on an intensive, customized preprocessing workflow that requires some reengineering when applied to other taxa. However, if the underlying principles and feature representation are retained, the model can be adapted straightforwardly by substituting or expanding the reference training set. While not a substitute for classical taxonomy (which requires full genomes and sometimes biological characterization), this approach rapidly highlights candidates warranting deeper investigation. Future work should focus on iterative retraining using newly discovered sequences and on including more phylogenetically diverse references.

## Supplementary data

Supplementary data are available online.

## Acknowledgements

We thank Nataša Mehle for leading the TOBAMO research project (ARIS L4-3179) that enabled the work presented in this paper.

## Data availability

Code, scripts, and additional supplementary materials supporting this study are deposited across three public GitHub repositories and permanently archived on Zenodo:

- **tobamo-analysis**: Contains dataset discovery routines (Serratus PalmID query yielding 253 candidate SRRs), manual curation workflows, clustering, phylogenetic placement scripts, and machine-learning classification models, https://github.com/nezapajek/tobamo-analysis, DOI: 10.5281/zenodo.23366618 (Pajek Arambašič et al., 2026a).
- **tobamo-snakemake**: Automated Snakemake assembly and candidate contig discovery pipeline covering read quality control, *de novo* assembly, similarity searches, and preliminary taxonomic assignment, https://github.com/nezapajek/tobamo-snakemake, DOI: 10.5281/zenodo.23366548 (Pajek Arambašič et al., 2026b).
- **tobamo-supp-data**: Large, additional supplementary materials: Supplementary Tables S3-S6 and Supplementary Files S1-S4 (multiple sequence alignments), h t t p s : / / g i t h u b. c o m / n e z a p a j e k / t o b a m o-s u p p-d a t a, DOI: 10.5281/zenodo.23366154 (Pajek Arambašič et al., 2026c).

## Funding

This work was supported by the project grant L4-3179 (Discovery and water-linked epidemiology of emergent tobamoviruses infecting crops) financed by the Slovenian Research and Innovation Agency (ARIS), the Ministry of Agriculture, Forestry and Food of the Republic of Slovenia, and BIA d.o.o. The work was also supported by the project grant GC-0001 (Artificial Intelligence for Science) and the core funding P4-0165 (Biotechnology and Plant Systems Biology) and P2-0209 (Artificial Intelligence and Intelligent Systems), all funded by ARIS.

## Ethics statement

No ethical approval or patient consent was required, as this study involved secondary analysis of publicly available data.

## Competing interests

No competing interests are declared.

## Author contributions statement

D.K., T.C., and K.B. conceived the analysis. N.P.A., T.C., D.K., K.B., L.K., and L.V. conducted the analysis. N.P.A., D.K., K.B., L.V., T.C., and L.K. wrote the manuscript. All authors have read, contributed to, and approved the final version of the manuscript.

## Notes

https://doi.org/10.5281/zenodo.23366618

https://doi.org/10.5281/zenodo.23366548

https://doi.org/10.5281/zenodo.23366154

### Competing Interest Statement

The authors have declared no competing interest.

