## Supplementary Materials for "Data mining of global sequence datasets markedly expands known diversity within the Tobamovirus genus"

### List of Supplementary Materials

**Supplementary Methods** Detailed description of the local tpdb2 and NCBI `nr` database setup for Snakemake pipeline, and description of machine learning framework for automated curation of putative tobamoviral contigs.

**Supplementary Results** Detailed description of the machine learning evaluation results.

**Supplementary Tables and Figures**

**Supplementary Table S1** Pairwise alignment metrics between ORFs and reference protein sequences.

**Supplementary Table S2** Feature importance rankings for the tobamovirus ORF classifier (Random Forest).

**Supplementary Tables S3-S6** are available at Zenodo.org [DOI: 10.5281/zenodo.23366154] and on GitHub `https://github.com/nezapajek/tobamo-supp-data`:

- **Table S3:** Serratus PalmID results from the initial query of 35 tobamoviral palmprints.
- **Table S4:** Metadata for 253 selected SRA accessions.
- **Table S5:** 510 selected contigs, including sequences, ORF breakdowns, model predictions, ground truths, and metadata.
- **Table S6:** Reference database used for model training.

**Supplementary Files S1-S4** are available at Zenodo.org [DOI: 10.5281/zenodo.23366154] and on GitHub `https://github.com/nezapajek/tobamo-supp-data`. Trimmed alignment of cluster representatives, reference sequences and outgroup for maximum-likelihood tree inference from:

- RdRp ORF1 (RdRp-ORF1_tree_alignment.fasta),
- RdRp ORF2 (RdRp-ORF2_tree_alignment.fasta),
- MP (MP_tree_alignment.fasta),
- CP (CP_tree_alignment.fasta).

**Supplementary Figure S1** Validation of contig length sampling distribution.

**Supplementary Figure S2** Feature enrichment analysis of the top 40 features in the ORF classifier.

**Supplementary Figure S3** Model performance comparison across histogram bin configurations

**Supplementary Figure S4** Threshold optimization analysis and performance metrics for the contig classifier.

**Supplementary Figure S5** Prediction-probability distributions at ORF and contig level

**Supplementary Figures S6-S9** Maximum likelihood phylogenetic trees of novel tobamoviral sequences inferred from RdRp ORF1, RdRp ORF2, MP, and CP, respectively.

S1.2.6 External evaluation on manually curated data and representative non-virga sequences . . 5

**S2 Supplementary Results: Detailed description of the machine learning evaluation results.** **6**

**S3 Supplementary Tables and Figures** **7**

### S1 Supplementary Methods

#### S1.1 Snakemake pipeline - local database setup

**Automated assembly and candidate contig discovery pipeline.** Short-read SRA accessions selected via Serratus PalmID (Edgar et al., 2022) were processed using an automated Snakemake workflow (v7.32.4) (Mölder et al., 2021). Raw sequencing reads were quality-trimmed using Trimmomatic (v0.39) (Bolger et al., 2014) and assembled *de novo* into contigs using SPAdes (v3.15.4) (Bankevich et al., 2012) or MEGAHIT (v1.2.9) (Li et al., 2015). Assembled contigs were length-filtered to retain sequences between 600 and 8,000 nt. Candidate contigs were searched sequentially using DIAMOND blastx (v0.9.14) (Buchfink et al., 2021) against:

1. A curated tobamoviral protein database (`tpdb2`, 2,177 protein sequences downloaded December 23, 2021, and compiled into a DIAMOND index on January 4, 2022).
2. A local build of the NCBI non-redundant protein database (`nr`, built on April 15, 2022).

Taxonomic assignment was performed using MEGAN6 (v6.21.7) (Beier et al., 2017) via the Lowest Common Ancestor (LCA) algorithm.

**NCBI `nr` Local Indexing & Version Tracking.** The local DIAMOND `nr` database index (`nr.dmnd`) is built from NCBI `nr` database releases using the `update_blastdb.pl` → `blastdbcmd` → `diamond makedb` procedure documented in the GitHub repository (`https://github.com/nezapajek/tobamo-snakemake`). This specific index is not distributed alongside the pipeline: NCBI `nr` is an unversioned, continuously growing snapshot with no archived releases, and a local build exceeds 100 GB, well beyond what repositories such as Zenodo or GitHub are intended to host. Re-indexing against a current `nr` release will incorporate newly deposited sequences, but candidate selection and the final machine learning classification are unaffected, because both are architecturally independent of the `nr` search. Only the `nr`-derived MEGAN taxonomic annotations can change.

#### S1.2 Detailed description of the machine learning pipeline for automated curation of putative tobamoviral contigs

This section provides a detailed description of the individual steps composing the machine learning pipeline used for automated curation of putative tobamoviral contigs. Each step is briefly summarized in the main text and described here in greater detail, following the same logical order.

##### S1.2.1 Reference sequence selection and creation of the training set.

**Obtaining references and pairwise alignment.** To construct a reference dataset suitable for supervised learning, we collected all NCBI whole-genome sequences assigned to genera within the *Virgaviridae* family (accessed January 20, 2025). These sequences were used to represent both the target category ( *Tobamovirus*) and closely related outgroup viruses from the family *Virgaviridae*. For each genus, pairwise sequence alignments were computed using MAFFT (v7.520) (Katoh and Standley, 2013).

**Clustering and selecting representatives.** Pairwise identity scores derived from these alignments were used to perform hierarchical clustering with single linkage using SciPy (v1.14.1). In the case of multipartite genomes, only the longest segment (RNA1) was used for selecting representatives. Clusters were defined using a 90% identity threshold. To limit representation bias arising from uneven sampling across species, only one representative sequence was selected per cluster. The final reference database consisted of 61 tobamoviral sequences and 27 outgroup sequences. For multipartite outgroup viruses, all genomic segments corresponding to the selected representative were included.

**Sequence augmentation and synthetic contig generation.** To increase the diversity of training examples and reflect realistic assembly conditions, we generated a synthetic dataset through sequence augmentation. From each genome in the reference database, 30 contigs were randomly sampled. Contig lengths were drawn from the empirical length distribution observed in the 510 identified tobamoviral candidate contigs. Specifically, we fitted a probability density curve to the observed lengths and used this model to sample the lengths for the synthetic contigs. To ensure the authenticity of the augmentation, we compared the fitted distribution of the original data against the distribution of the newly sampled lengths, as shown in Supplementary Figure S1.

For multipartite genomes, the genomic segment to sample from was selected in proportion to its relative length within the complete genome. A contig of the chosen length was then sampled from that segment. This strategy ensured that the augmented dataset closely matched the length variability and fragmentation patterns of real contigs. Each synthetic contig was labeled according to its source genome as either tobamoviral or outgroup.

##### S1.2.2 Sequence alignment with reference ORFs, alignment, and sequence-based features

To construct a comprehensive feature set for contig classification, we derived both alignment-based and sequence-based features from each predicted ORF. These features capture evolutionary similarity, sequence conservation, and basic structural characteristics, serving as the input for downstream machine learning models. The following subsections describe the procedures for ORF extraction, pairwise alignment with reference sequences, feature aggregation, and the composition of the final dataset.

**ORF extraction.** We extracted ORFs using orfipy (Singh and Wurtele, 2021) with the options `-min 300 -between-stops`, which allows ORFs without START or STOP codon. In addition, we used biopython (Cock et al., 2009) `Bio.Seq.` module's `translate()` function in all 6 reading frames, to detect all coding sequences without START and STOP codons.

**Pairwise alignment.** The next step involved pairwise alignment of all ORFs against all reference dataset amino acid sequences, excluding those from their parent genome to prevent overfitting. Although four viral proteins were initially considered (RdRp ORF1, RdRp ORF2, coat protein (CP), and movement protein (MP)), only three were retained for downstream analyses. The MP was excluded due to a lack of homologous representation across all *Virgaviridae* members, as movement functions are not uniformly encoded by a single, comparable ORF, which limits cross-group alignment and comparability. We used MAFFT (v.7.520) (Katoh and Standley, 2013) for pairwise sequence alignment. For each alignment, we calculated multiple alignment quality metrics: identity score, M, N, alignment length, N/alignment length ratio, aln_orf_len, gap openings, and gap ratio. Detailed descriptions of these metrics are provided in the Supplementary table **??**.

**Alignment-based features.** Following pairwise alignment computation, we performed a comprehensive aggregation step to summarize alignment statistics for downstream classification. For each ORF, we calculated the minimum, mean, and maximum values of each alignment metric across all reference sequences. This aggregation was performed for each combination of protein type (RdRp ORF1, RdRp ORF2, and coat protein CP) and reference group (tobamoviruses and outgroup viruses), resulting in a feature matrix where each contig was represented by 3 proteins $\times$ 2 reference groups $\times$ 8 metrics $\times$ 3 aggregation statistics = 144 alignment-derived features. This aggregation strategy captured both the best matches (minimum/maximum values) and overall similarity patterns (mean values) between query sequences and reference taxonomic groups, thereby providing robust statistical summaries for machine-learning classification.

**Sequence-based features.** In addition to the alignment-derived features, we incorporated four sequence-based features for each ORF: (1) ORF length (orf_len), representing the number of amino acids in the predicted open reading frame, (2) contig length (contig_length), indicating the total nucleotide length of the parent contig, (3) stop codon count (stop_codons), quantifying the number of premature stop codons within the ORF sequence as an indicator of sequence quality, and (4) ORF prediction method (orf_acc_method), a binary feature distinguishing between ORFs identified using ORFipy versus those obtained through six-frame translation. These additional features provided complementary insights into sequence characteristics and the prediction methodology.

**Dataset for machine learning.** The final feature set comprised 148 features per ORF (144 alignment-based and 4 sequence-based). In total, the dataset contained 34,492 ORFs, of which 29,311 were forward-strand and 5,181 reverse-strand sequences. Model training was performed using only forward-strand ORFs, excluding those RNA segments encoding only movement protein (MP) or triple gene block (TGB) proteins - typically RNA3, although not all RNA3 segments. The exact inclusion or exclusion of sequences is detailed in Supplementary Table S6 (column "training"). For testing, we used ORFs from all genomic segments and from reverse strands. This setup enabled the model to learn from biologically representative forward-strand coding regions while evaluating its generalization to unseen genomic segments and reverse-strand sequences, thereby enhancing both robustness and realism of the classification task. With this dataset assembled, we trained and compared multiple machine learning classifiers to identify the most effective model for distinguishing tobamoviral and non-tobamoviral ORFs.

##### S1.2.3 Model selection for ORF classifier

Model selection involves testing different algorithms to identify which best learns the relationship between input features and their correct classification labels. Accordingly, we compared several commonly used machine learning classifiers to determine the most suitable approach for classifying ORFs into tobamovirus or outgroup, based on their feature representations.

We initially evaluated Logistic Regression, Support Vector Machines (SVM), Decision Trees, Naïve Bayes, K-Nearest Neighbors (KNN), and Random Forest, implemented using the scikit-learn library (Pedregosa et al., 2011). Each model underwent hyperparameter tuning via grid search, optimizing parameters such as regularization strength for logistic regression, kernel types for SVM, and the number of estimators for Random Forest.

Model selection was performed using a repeated stratified cross-validation framework (5 iterations $\times$ 5 folds).

Within each outer fold, candidate classifiers were tuned using an inner 5-fold grid search (GridSearchCV, scoring=accuracy) on the training partition, after preprocessing and feature scaling. The best-performing model from the inner cross-validation was then evaluated on the corresponding held-out outer test fold. Final model selection was based on the highest mean outer-fold test accuracy across all iterations, resulting in a Random Forest classifier (200 estimators, `maximum depths: 50`) used to predict whether a given ORF belongs to tobamovirus or the outgroup.

For feature interpretation, we scored and ranked feature importance using the Random Forest ORF classifier trained on the reference training dataset, quantified by scikit-learn's mean decrease in impurity (MDI) importance score.

Feature enrichment analysis was performed on MDI importance values from the final Random Forest model, in which features were ranked by importance, and the top 40 of the 148 total features were used as the target set. Each feature was assigned to categories by parsing feature names, including source group (tobamovirus, outgroup, or generic/basic), protein group (ORF1, ORF2, CP), and metric family (*e.g.*, *identity_ score*, *gap_ ratio*, *gap_ openings*, *aln_ len*, *aln_ orf_ len*, and *N/aln_ len*). Enrichment was tested separately for metric-based and protein-based category schemes using one-sided hypergeometric tests. For each category, a one-sided hypergeometric test gave the probability of observing at least as many of its features among the top 40 as were found. The expected count was the number of top-ranked features multiplied by the category's share of all features, and the enrichment factor was the ratio of the observed to the expected count.

##### S1.2.4 Contig classifier, aggregation strategies of ORF-level predictions using stacked generalization

Each contig may include a varying number of individual ORF predictions. Because biological interpretation requires a single prediction per contig, ORF-level predictions were aggregated into a contig-level classification using stacking (stacked generalization as introduced in Wolpert (1992)).

Two aggregation strategies were evaluated: (i) **Extreme prediction**, which assigns the contig label according to the ORF with the most confident probability (furthest from 0.5), and (ii) **Prediction binning** using a logistic regression meta-classifier applied to a vector of binned ORF probabilities (10 bins). We refer to the contig classifier using extreme prediction aggregation as Extreme, and the contig classifier using prediction binning as Binned. Both aggregation strategies were compared using cross-validated performance metrics and calibration analyses.

**Extreme prediction.** In the first approach, the prediction for the entire contig was determined by selecting the single ORF with the most extreme probability score, that is, the score farthest from 0.5. This method assumes that the most confident ORF-level prediction provides the strongest evidence for the contig classification. We refer to the contig classifier using extreme prediction aggregation as Extreme.

**Prediction binning.** In the second approach, we constructed a histogram of ORF probability scores using ten equally spaced bins and used this distribution as input to a logistic regression classifier, which we later refer to as Binned. Unlike the Extreme prediction, the prediction binning strategy incorporates the overall pattern of predictions across all ORFs within a contig, thereby capturing more comprehensive evidence rather than relying on a single extreme value. We evaluated several bin sizes (5, 10, 15, 20), and found that 10 bins provided a good balance between resolution and performance, as shown in Supplementary Figure S3.

Performance metrics, including accuracy, F1 score, precision, recall, and AUC, were compared between the two contig classifiers, and Binned was selected as the best-performing approach. We also performed a threshold optimization analysis across 150 cross-validation folds to determine the optimal threshold, see Supplementary Figure S4. The final machine learning prediction for each contig was a single label indicating whether it was tobamoviral or non-tobamoviral.

##### S1.2.5 Evaluation of the contig classifier

We conducted a comprehensive evaluation to compare contig-level prediction strategies and assess model robustness. A rigorous cross-validation framework was implemented, consisting of 30 independent iterations, each using 5-fold stratified cross-validation, for a total of 150 evaluation rounds. This extensive testing protocol was designed to provide statistically robust performance estimates and minimize the impact of random variation in data splits. Within each fold, a leave-one-virus-out strategy was applied for ORF-level predictions. Specifically, when a particular virus species was designated as the test set, all remaining virus species formed the training set for the Random Forest ORF classifier. This approach ensured that ORF-level predictions were made without direct exposure to sequences from the same viral species, thereby providing a rigorous evaluation of the model's generalizability. Performance was evaluated using multiple metrics, including accuracy, F1 score, precision, recall, and area under the ROC curve (AUC). Statistical significance was assessed using 95% confidence intervals calculated from 30 iterations.

**Calibration analysis.** We evaluated the probability calibration using reliability diagrams at the contig level for both classifiers (Binned and Extreme). For each method and each cross-validation iteration (pooling predictions across the five folds), we computed binned calibration points (`prob_true`, `prob_pred`) with `sklearn.calibration.calibration_curve` using 10 bins from `ground_truth` and the predicted probability (`prob_1`). Iteration-specific curves were then interpolated to a common 0–1 grid (100 points) and summarized as mean $\pm$ SD, with the identity line ($y = x$) used as the perfect-calibration reference.

**Threshold analysis.** We performed a post hoc threshold analysis on the cross-validation predictions of the selected Binned classifier. In each of the 150 iteration–fold units (30 iterations $\times$ 5 folds), the decision threshold was tuned for accuracy by inner 10-fold cross-validation on the training data and applied to the held-out fold. We summarized the variability of the tuned thresholds between and within iterations and compared held-out accuracy at the tuned threshold with accuracy at the default threshold of 0.5. To check the robustness, we also applied a single global threshold to all folds, from 0.30 to 0.70 in steps of 0.01. Because tuning did not improve accuracy (mean change of $-0.0004$), the final model uses the default threshold of 0.5.

##### S1.2.6 External evaluation on manually curated data and representative non-virga sequences

We tested the model beyond the training data in two ways to evaluate its robustness under realistic and challenging conditions, and to observe how the model behaves when encountering real-life data and how well it mirrors expert human judgment when applied to real candidate contigs.

**Viral representatives (non-*Virgaviridae* test).** To test the model's ability to distinguish tobamovirus sequences from more divergent viral sequences, we downloaded all NCBI viral representatives (accessed 2025-01-16) and duplicated them at the genus level, generating 1,398 representatives. These were screened with DIAMOND against our local tobamoviral database, and 41 sequences (2.93% of the representatives) passed the DIAMOND filter and were retained as candidates for model evaluation.

**Test on manually curated data.** In addition, we applied the model to 510 discovered contigs in this study and compared the predictions with manually curated ground truth provided by domain scientists. For each contig, we examined the predicted label in the context of its assigned category, assessing agreement between model predictions and manually determined expert annotations for tobamovirus-like contigs, other viral groups, and misassembled sequences. This per-category evaluation provided insight into which contig types were most accurately classified and highlighted any systematic discrepancies between model predictions and manual curation.

### S2 Supplementary Results: Detailed description of the machine learning evaluation results.

#### Machine learning approach aids in data curation step

In the first step, we performed ORF-level classification and tested several classifiers for this task. Among the classifiers tested, Random Forest consistently demonstrated superior performance, being selected in 22 out of 25 cross-validation folds (88%) with a mean accuracy of 0.7671 ± 0.0215, significantly outperforming the second-best algorithm, SVM, which was selected in only 3 folds (12%) with a mean accuracy of 0.7524 ± 0.0144. Among the Random Forest hyperparameter combinations tested, the best single performance was achieved with `max_depth: 40` and `n_estimators: 200` (accuracy 0.7953), but we selected the more robust combination of `max_depth: 50` and `n_estimators: 200` for subsequent analyses. This configuration demonstrated superior stability, being selected in 4 of the 22 Random Forest folds, with a mean accuracy of 0.7799 ± 0.0085, a mean AUC of 0.6391 ± 0.0061, and a mean F1 score of 0.8624 ± 0.0069. The selection prioritized consistent performance over peak single-fold results, ensuring model reliability across different data partitions.

A *post hoc* check found that three of the six model-selection classifiers had not been run with a fixed random seed (only Random Forest was). Rerunning model selection with a fixed seed changed results negligibly (0.7643 ± 0.0212 *vs.* 0.7671 ± 0.0215 originally). The evaluation stage was already fully seeded but had only been tested at one seed. Rerunning it at two additional seeds gave accuracies of 0.9728 and 0.9734, both within the originally reported 95% CI (0.9727–0.9743), confirming the results are not seed-dependent.

Feature-importance analysis of the final Random Forest ORF classifier suggests the absence of a single dominant feature. Of 148 features, the top 10 and top 20 accounted for 16.4% and 28.8% of total importance, respectively. The most influential features were primarily alignment-derived metrics (identity, gap, and alignment-length descriptors) and the number of stop codons, which gives information about the strand and the correct open reading frame. Importance was balanced between tobamovirus- and outgroup-derived features (46.8% *vs.* 49.4%), and was similarly distributed across ORF1, ORF2, and CP feature groups (33.9%, 33.2%, and 29.1%) (see Supplementary table S2). Hypergeometric enrichment analysis of the top 40 features identified significant overrepresentation of *identity_ score* and *gap_ ratio* descriptors (14/18 and 13/18 features, enrichment 2.88x and 2.67x, p = 2.15e-06 and 2.39e-05, both Bonferroni-significant), whereas protein-category enrichments were not significant (all p > 0.05). The top 40 features jointly explained 47.7% of total feature importance (see Supplementary Figure S2).

In the second step, we performed contig-level classifications, aggregating ORF-level classifications for each contig when multiple ORFs were detected. Across 30 independent runs of five-fold cross-validation, the Binned classifier consistently outperformed the Extreme classifier, see Figure 2 (a,b) in the main text. The Binned contig classifier achieved an average accuracy of 0.9735 (95% CI: 0.9727–0.9743) with an F1 score of 0.9806 (95% CI: 0.9801–0.9812). Precision (0.9925) and recall (0.9690) were both high, and discrimination was nearly perfect, with an AUC of 0.9971 (95% CI: 0.9969–0.9972). By contrast, the Extreme classifier reached an accuracy of 0.9231 (95% CI: 0.9217–0.9246) and an F1 score of 0.9475 (95% CI: 0.9465–0.9484), with notably lower precision (0.9003) despite a recall close to 1.0. Calibration analysis further favored the Binned classifier, whose predicted probabilities closely matched the observed outcome frequencies, indicating near-perfect calibration. This conclusion is supported by the threshold-optimization analysis, which showed that additional threshold tuning was not necessary, see Supplementary Figure S4.

### S3 Supplementary Tables and Figures

Supplementary Table S1: Pairwise alignment metrics calculated between an ORF and a reference protein sequence.

| Metric | Definition | Calculation |
|---|---|---|
| Identity Score ($identity\_score$) | Proportion of mismatched positions relative to aligned positions. | $1 - \frac{M}{N}$, where $M$ = mismatches and $N$ = aligned positions. |
| Mismatches ($M$) | Number of positions with different residues (excluding gaps). | Count positions where $seq1[i] \neq seq2[i]$ and neither is a gap. |
| Aligned Positions ($N$) | Positions where both sequences contain residues (no gaps). | Count positions where $seq1[i] \neq -$ and $seq2[i] \neq -$. |
| Alignment Length ($aln\_len$) | Span of the alignment region. | $aln\_len = last\_match - first\_match + 1$ |
| ORF Length ($orf\_len$) | Length of ORF sequence excluding gaps. | Count non-gap residues in ORF sequence. |
| Reference Length ($ref\_len$) | Length of reference sequence excluding gaps. | Count non-gap residues in reference sequence. |
| Alignment Density ($N/aln\_len$) | Ratio of aligned positions to alignment length. | $N/aln\_len$ |
| Gap Openings ($gap\_openings$) | Number of transitions from residue to gap. | Count positions where a residue is followed by a gap (-). Sum for both sequences, excluding terminal gaps. |
| Gap Ratio ($gap\_ratio$) | Ratio of gap openings to alignment length. | $gap\_openings/aln\_len$ |
| ORF Coverage in Alignment ($aln\_orf\_len$) | Portion of ORF included in the alignment. | $aln\_len-$(gaps in ORF sequence within alignment region) |

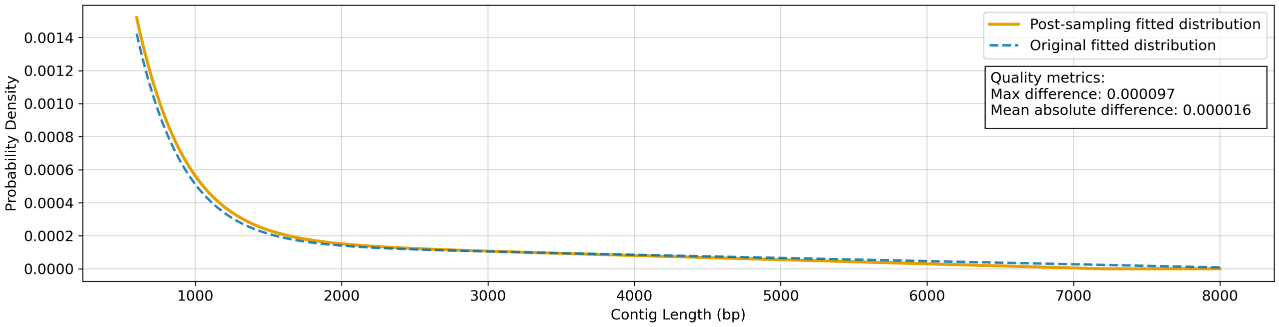

Supplementary Figure S1: **Validation of contig length sampling distribution.** This figure compares two fitted contig-length distributions across 600-8000 bp. The blue dashed curve is the original target distribution loaded from the precomputed length-frequency model, and the orange solid curve is a new fit to the sampled contigs FASTA used for downstream training. Both curves use the same functional form, $f(x) = ae^{-bx} + cx + d$, and are normalized before comparison. The close overlap, supported by low maximum and mean absolute differences reported in the annotation box, indicates that the sampling process preserved the intended length profile.

**Alt text:** Line chart of probability density versus contig length from six hundred to eight thousand base pairs with two nearly overlapping curves: orange solid for post-sampling fitted distribution and blue dashed for original fitted distribution, plus a small box reporting max and mean absolute differences.

Supplementary Table S2: Feature importance values for tobamovirus ORF classifier (Random Forest classifier), the final random forest model, ranked from highest to lowest importance. Features represent various alignment metrics between contigs and reference sequences for different viral proteins (RdRp ORF1, RdRp ORF2, CP), comparing tobamovirus and outgroup references.

| Rank | Feature | Importance |
|---|---|---|
| 1 | tobamo_orf1_identity_score_max | 0.0217 |
| 2 | outgroup_orf1_identity_score_max | 0.0216 |
| 3 | outgroup_orf2_identity_score_max | 0.0163 |
| 4 | stop_codons | 0.0163 |
| 5 | outgroup_orf2_identity_score_mean | 0.0155 |
| 6 | outgroup_orf2_identity_score_min | 0.0151 |
| 7 | outgroup_orf1_N/aln_len_max | 0.0149 |
| 8 | tobamo_orf1_N/aln_len_max | 0.0145 |
| 9 | tobamo_orf1_gap_openings_min | 0.0145 |
| 10 | outgroup_orf2_gap_ratio_mean | 0.0134 |
| 11 | tobamo_orf2_identity_score_max | 0.0132 |
| 12 | contig_length | 0.0129 |
| 13 | outgroup_orf1_gap_ratio_min | 0.0128 |
| 14 | outgroup_cp_identity_score_max | 0.0128 |
| 15 | tobamo_orf1_gap_ratio_min | 0.0127 |
| 16 | outgroup_orf1_gap_openings_min | 0.0123 |
| 17 | outgroup_cp_gap_ratio_min | 0.0122 |
| 18 | outgroup_orf1_identity_score_mean | 0.0119 |
| 19 | tobamo_orf1_gap_ratio_mean | 0.0118 |
| 20 | tobamo_orf2_identity_score_mean | 0.0114 |
| 21 | outgroup_orf2_gap_ratio_min | 0.0113 |
| 22 | tobamo_orf2_identity_score_min | 0.0110 |
| 23 | outgroup_orf2_gap_ratio_max | 0.0110 |
| 24 | tobamo_orf1_identity_score_mean | 0.0107 |
| 25 | outgroup_cp_N/aln_len_max | 0.0103 |
| 26 | outgroup_orf2_N/aln_len_max | 0.0102 |
| 27 | tobamo_orf2_gap_ratio_max | 0.0098 |
| 28 | tobamo_orf2_gap_ratio_min | 0.0097 |
| 29 | tobamo_cp_identity_score_max | 0.0097 |
| 30 | tobamo_cp_gap_ratio_min | 0.0096 |
| 31 | tobamo_cp_identity_score_mean | 0.0093 |
| 32 | outgroup_orf1_aln_len_mean | 0.0090 |
| 33 | tobamo_cp_gap_ratio_mean | 0.0090 |
| 34 | outgroup_orf1_aln_len_min | 0.0089 |
| 35 | tobamo_orf2_gap_ratio_mean | 0.0087 |
| 36 | tobamo_orf2_N/aln_len_max | 0.0085 |
| 37 | tobamo_cp_gap_openings_mean | 0.0083 |
| 38 | tobamo_orf1_gap_openings_mean | 0.0082 |
| 39 | tobamo_orf1_identity_score_min | 0.0082 |
| 40 | outgroup_cp_gap_ratio_mean | 0.0078 |
| 41 | tobamo_cp_M_mean | 0.0077 |
| 42 | outgroup_cp_gap_openings_mean | 0.0073 |
| 43 | outgroup_cp_identity_score_mean | 0.0073 |
| 44 | tobamo_cp_N/aln_len_max | 0.0072 |
| 45 | tobamo_orf1_gap_ratio_max | 0.0072 |
| 46 | tobamo_orf2_gap_openings_mean | 0.0072 |
| 47 | outgroup_orf1_identity_score_min | 0.0071 |
| 48 | tobamo_cp_identity_score_min | 0.0071 |
| 49 | outgroup_cp_gap_openings_min | 0.0071 |
| 50 | outgroup_cp_M_min | 0.0070 |
| 51 | outgroup_cp_gap_ratio_max | 0.0069 |
| 52 | outgroup_orf2_gap_openings_mean | 0.0069 |
| 53 | outgroup_orf1_gap_ratio_mean | 0.0069 |

Continued on next page

**Supplementary Table S2 – continued from previous page**

| Rank | Feature | Importance |
|---|---|---|
| 54 | tobamo_orf2_gap_openings_min | 0.0068 |
| 55 | outgroup_cp_M_mean | 0.0067 |
| 56 | tobamo_orf1_aln_len_mean | 0.0067 |
| 57 | outgroup_orf1_gap_openings_mean | 0.0065 |
| 58 | outgroup_orf1_gap_ratio_max | 0.0064 |
| 59 | outgroup_orf2_M_min | 0.0064 |
| 60 | outgroup_cp_identity_score_min | 0.0064 |
| 61 | tobamo_orf2_M_min | 0.0063 |
| 62 | outgroup_orf1_N/aln_len_mean | 0.0062 |
| 63 | tobamo_orf1_aln_len_min | 0.0062 |
| 64 | tobamo_cp_gap_ratio_max | 0.0061 |
| 65 | outgroup_orf2_N/aln_len_mean | 0.0061 |
| 66 | tobamo_orf2_N/aln_len_mean | 0.0060 |
| 67 | outgroup_orf1_aln_len_max | 0.0059 |
| 68 | outgroup_orf2_N_min | 0.0058 |
| 69 | outgroup_cp_N/aln_len_mean | 0.0057 |
| 70 | tobamo_cp_N/aln_len_min | 0.0056 |
| 71 | outgroup_orf2_aln_len_max | 0.0056 |
| 72 | outgroup_orf2_aln_len_min | 0.0056 |
| 73 | tobamo_cp_aln_len_min | 0.0056 |
| 74 | tobamo_cp_N_mean | 0.0056 |
| 75 | tobamo_orf2_aln_len_mean | 0.0056 |
| 76 | tobamo_cp_aln_orf_len_min | 0.0055 |
| 77 | orf_len | 0.0055 |
| 78 | outgroup_orf2_gap_openings_min | 0.0055 |
| 79 | outgroup_orf2_aln_len_mean | 0.0055 |
| 80 | tobamo_orf2_N/aln_len_min | 0.0055 |
| 81 | tobamo_orf1_N/aln_len_mean | 0.0055 |
| 82 | outgroup_cp_N_mean | 0.0055 |
| 83 | tobamo_cp_M_max | 0.0054 |
| 84 | tobamo_orf1_aln_len_max | 0.0053 |
| 85 | outgroup_orf2_N/aln_len_min | 0.0053 |
| 86 | outgroup_orf2_M_mean | 0.0053 |
| 87 | outgroup_cp_aln_orf_len_min | 0.0053 |
| 88 | outgroup_cp_M_max | 0.0052 |
| 89 | tobamo_cp_N/aln_len_mean | 0.0051 |
| 90 | outgroup_cp_N/aln_len_min | 0.0050 |
| 91 | tobamo_cp_M_min | 0.0050 |
| 92 | outgroup_orf1_gap_openings_max | 0.0049 |
| 93 | tobamo_orf2_gap_openings_max | 0.0049 |
| 94 | outgroup_cp_aln_len_min | 0.0049 |
| 95 | tobamo_orf2_aln_len_min | 0.0049 |
| 96 | tobamo_orf2_M_mean | 0.0048 |
| 97 | tobamo_orf2_N_mean | 0.0048 |
| 98 | tobamo_orf2_N_min | 0.0047 |
| 99 | outgroup_cp_N_max | 0.0047 |
| 100 | outgroup_orf1_N/aln_len_min | 0.0047 |
| 101 | tobamo_orf1_N/aln_len_min | 0.0047 |
| 102 | tobamo_orf1_M_min | 0.0046 |
| 103 | tobamo_cp_N_max | 0.0045 |
| 104 | tobamo_orf2_aln_len_max | 0.0044 |
| 105 | outgroup_cp_N_min | 0.0043 |
| 106 | tobamo_orf2_M_max | 0.0043 |
| 107 | outgroup_orf2_N_mean | 0.0043 |
| 108 | tobamo_cp_aln_orf_len_mean | 0.0042 |
| 109 | outgroup_orf2_aln_orf_len_min | 0.0041 |
| 110 | tobamo_cp_aln_len_mean | 0.0041 |
| 111 | outgroup_orf1_M_min | 0.0040 |

Continued on next page

**Supplementary Table S2 – continued from previous page**

| Rank | Feature | Importance |
|---|---|---|
| 112 | outgroup_orf2_M_max | 0.0040 |
| 113 | outgroup_orf1_M_mean | 0.0040 |
| 114 | outgroup_cp_aln_len_max | 0.0040 |
| 115 | tobamo_cp_N_min | 0.0039 |
| 116 | outgroup_cp_aln_orf_len_mean | 0.0038 |
| 117 | tobamo_orf2_N_max | 0.0038 |
| 118 | outgroup_orf2_gap_openings_max | 0.0037 |
| 119 | tobamo_orf2_aln_orf_len_min | 0.0037 |
| 120 | tobamo_orf1_gap_openings_max | 0.0037 |
| 121 | outgroup_cp_aln_len_mean | 0.0036 |
| 122 | tobamo_orf2_aln_orf_len_mean | 0.0035 |
| 123 | tobamo_orf1_N_min | 0.0035 |
| 124 | tobamo_orf1_aln_orf_len_min | 0.0035 |
| 125 | tobamo_cp_aln_len_max | 0.0035 |
| 126 | outgroup_orf1_N_min | 0.0035 |
| 127 | outgroup_orf2_N_max | 0.0033 |
| 128 | tobamo_orf1_M_mean | 0.0033 |
| 129 | tobamo_cp_gap_openings_max | 0.0033 |
| 130 | tobamo_cp_gap_openings_min | 0.0033 |
| 131 | outgroup_cp_gap_openings_max | 0.0032 |
| 132 | outgroup_orf1_M_max | 0.0032 |
| 133 | outgroup_orf2_aln_orf_len_mean | 0.0032 |
| 134 | outgroup_orf1_N_mean | 0.0032 |
| 135 | tobamo_orf1_M_max | 0.0032 |
| 136 | tobamo_orf1_aln_orf_len_mean | 0.0032 |
| 137 | tobamo_orf1_N_mean | 0.0030 |
| 138 | outgroup_orf1_aln_orf_len_min | 0.0029 |
| 139 | outgroup_orf1_aln_orf_len_mean | 0.0028 |
| 140 | orf_acc_method | 0.0028 |
| 141 | tobamo_orf2_aln_orf_len_max | 0.0027 |
| 142 | outgroup_cp_aln_orf_len_max | 0.0026 |
| 143 | outgroup_orf2_aln_orf_len_max | 0.0026 |
| 144 | outgroup_orf1_N_max | 0.0025 |
| 145 | tobamo_orf1_aln_orf_len_max | 0.0025 |
| 146 | tobamo_cp_aln_orf_len_max | 0.0024 |
| 147 | tobamo_orf1_N_max | 0.0024 |
| 148 | outgroup_orf1_aln_orf_len_max | 0.0024 |

**Note:** Features are named using the pattern `[reference_type]_[protein]_[metric]_[aggregation]`, where reference_type is either "tobamo" (tobamovirus references) or "outgroup" (non-tobamovirus references), protein is "orf1", "orf2", or "cp" (coat protein), metric represents alignment statistics (*identity_score*, *gap_ratio*, *N/aln_len*, *etc.*), and aggregation indicates the statistical summary (*min*, *max*, *mean*) across all reference sequences of that type.

**a Metric categories - enrichment results**

| Category | Observed | Expected | Enrichment | P-value | Significant |
|---|---|---|---|---|---|
| **Identity score** | **14** | **4.9** | **2.88** | **2.15e-06** | **Yes** |
| **Gap ratio** | **13** | **4.9** | **2.67** | **2.39e-05** | **Yes** |
| N aln len ratio | 5 | 4.9 | 1.03 | 5.69e-01 | No |
| Gap openings | 4 | 4.9 | 0.82 | 7.75e-01 | No |
| Aln len | 2 | 4.9 | 0.41 | 9.80e-01 | No |
| Contig length | 1 | 0.3 | 3.70 | 2.70e-01 | No |
| Stop codons | 1 | 0.3 | 3.70 | 2.70e-01 | No |

**b Metric categories - frequency**

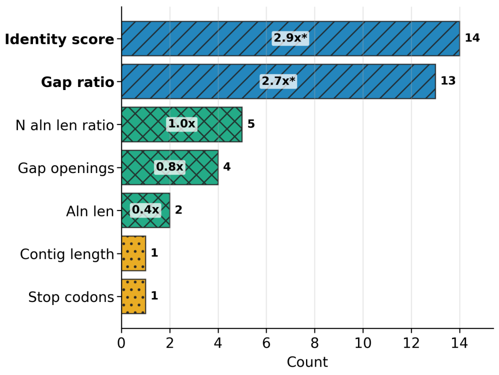

**c Protein categories - enrichment results**

| Category | Observed | Expected | Enrichment | P-value | Significant |
|---|---|---|---|---|---|
| Tobamo orf1 | 8 | 6.5 | 1.23 | 2.99e-01 | No |
| Outgroup orf1 | 7 | 6.5 | 1.08 | 4.87e-01 | No |
| Outgroup orf2 | 7 | 6.5 | 1.08 | 4.87e-01 | No |
| Tobamo orf2 | 7 | 6.5 | 1.08 | 4.87e-01 | No |
| Tobamo cp | 5 | 6.5 | 0.77 | 8.41e-01 | No |
| Outgroup cp | 4 | 6.5 | 0.62 | 9.39e-01 | No |
| Contig length | 1 | 0.3 | 3.70 | 2.70e-01 | No |
| Stop codons | 1 | 0.3 | 3.70 | 2.70e-01 | No |

**d Protein categories - frequency**

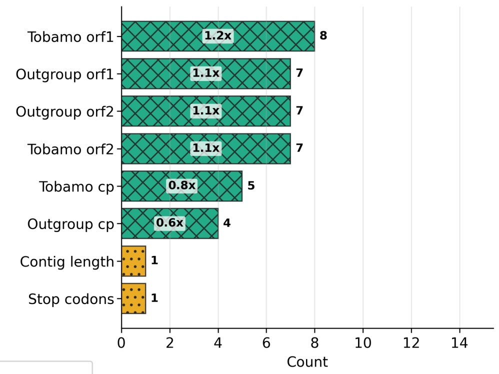

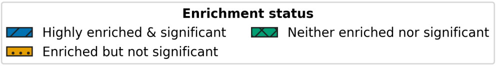

Supplementary Figure S2: **Feature category enrichment analysis and frequency distribution with enrichment status for the top 40 most important features in the tobamovirus contig classifier.** (a-c) Tables summarizing enrichment results for metric categories (a) and protein categories (c) with observed versus expected feature counts, enrichment factors, and significance values. (b,d) Plots showing the frequency distribution of top 40 features by metric categories (b) and protein categories (d) with enrichment factors displayed within bars and statistical significance indicated by asterisks (*p<0.05). Color and hatch coding and hatch patterns represent enrichment status: highly enriched and significant (blue, diagonal hatch), enriched but not significant (orange, filled dots hatch), and neither enriched nor significant (green, crossed diagonal grid hatch). The analysis identifies two significant metric categories, identity score and gap ratio, while no protein categories are statistically significant.

**Alt text (Panel a):** A table of enrichment results for metric categories, where identity score and gap ratio are statistically significant. **Alt text (Panel b):** Identity score and gap ratio have the highest feature counts, fourteen and thirteen, and are the only categories marked statistically significant, with the strongest enrichment. All other metric categories are lower, from five to one, and are not significant. **Alt text (Panel c):** A table of enrichment results for protein categories, where no metrics are statistically significant. **Alt text (Panel d):** Protein category counts are broadly distributed across tobamovirus and outgroup proteins, with tobamovirus open reading frame one highest at eight. Categories for open reading frame one and two cluster near seven, coat protein is lower, and no category is statistically significant.

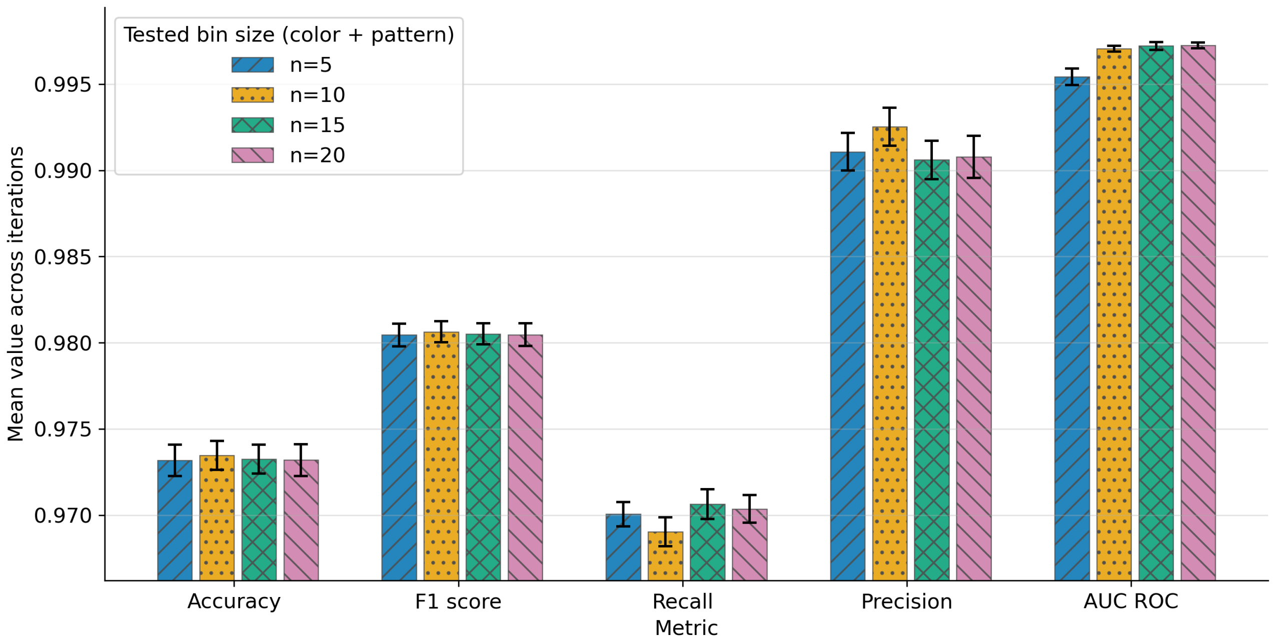

Supplementary Figure S3: **Model performance comparison across different bin configurations with 95% confidence intervals.** The grouped bar chart shows mean classification performance across five metrics: accuracy, F1 score, recall, precision, and AUROC. Within each metric group, bars correspond to tested histogram bin sizes ($n = 5$, 10, 15, 20), distinguished by both color and hatch pattern. Error bars show 95% confidence intervals estimated from repeated runs with different random seeds using a $t$-distribution. The $y$-axis is zoomed to the observed performance range to make small differences between configurations easier to compare. Overall, the ten-bin bar shows a slight performance edge, most clearly in precision and among the top values for accuracy and F1 score.

**Alt text:** Grouped bar chart comparing model performance across five metrics: accuracy, F one score, recall, precision, and area under the receiver operating characteristic curve. Each group evaluates four different histogram bin sizes: five, ten, fifteen, and twenty. All bars are high and close together, and each bar includes a narrow 95% confidence interval. The ten-bin bars are slightly higher overall, especially for precision.

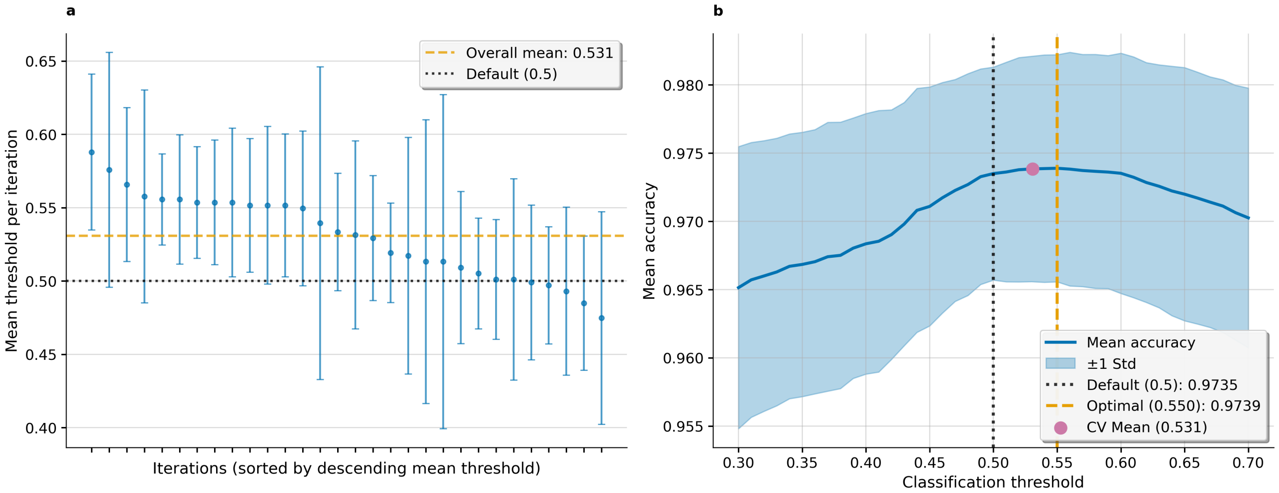

Supplementary Figure S4: **Threshold optimization analysis and performance relationship for the tobamovirus contig classifier.** (a) Distribution of accuracy improvements comparing optimized thresholds versus default threshold (p=0.5) across 150 cross-validation folds, showing minimal mean improvement (-0.0004) with most folds clustered around zero improvement. (b) Threshold-accuracy relationship curve demonstrating near-optimal performance at the default threshold, with the theoretical optimal threshold at 0.550 (accuracy = 0.9984) providing negligible improvement over default performance (accuracy = 0.9980). Cross-validation mean threshold (0.531) and confidence intervals (±1 std) indicate well-calibrated model probabilities. The analysis validates the use of the default threshold (p=0.5) for deployment, as the complexity of threshold optimization is not justified by the marginal performance gain of +0.0004 in accuracy.

**Alt text (Panel a):**Panel a plots threshold per iteration with error bars. Values cluster near zero point five three, decline slightly across iterations, and mostly overlap the zero point five reference. **Alt text (Panel b):** Panel b shows accuracy rising with threshold, peaking near zero point five five, then tapering slightly. A shaded uncertainty band surrounds the curve, with default and optimized markers.

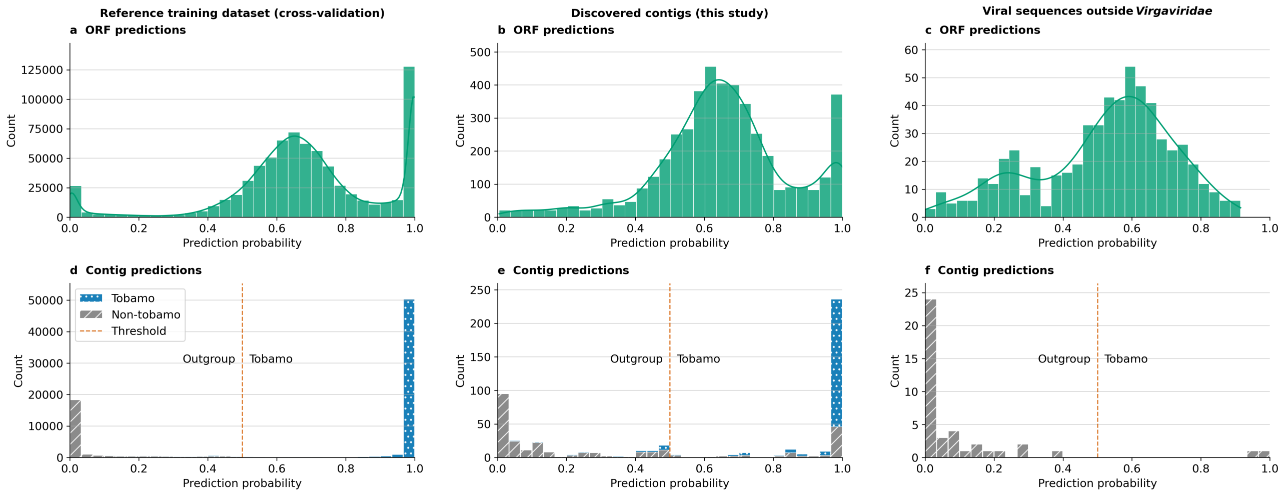

Supplementary Figure S5: Prediction-probability distributions at ORF level (a–c) and contig level (d–f) across three datasets: reference training data (cross-validation), discovered contigs from this study, and viral sequences outside *Virgaviridae*. ORF panels present probability histograms with density overlays, while contig panels show stacked category composition (Tobamo *vs.* Non-tobamo). The dashed line marks the decision threshold at $p = 0.5$.

**Alt text (Panel a):** Bimodal open reading frame probability distribution in reference cross-validation data, with broad mass around zero point six to zero point seven, a dominant spike near one point zero, and a smaller spike near zero.

**Alt text (Panel b):** Open reading frame probabilities for discovered contigs are mainly unimodal, peaking near zero point six to zero point seven, with secondary mass near one point zero and relatively few values below zero point four.

**Alt text (Panel c):** Open reading frame probabilities for viral sequences outside Virgaviridae show a broad peak near zero point six, a smaller shoulder around zero point two to zero point three, and tapering frequencies above zero point eight.

**Alt text (Panel d):** Contig-level reference cross-validation counts are concentrated at extremes, with Non-tobamo near zero and very large Tobamo near one, while very few contigs fall around the zero point five threshold.

**Alt text (Panel e):** Contig-level discovered-contig counts split toward both extremes, with Non-tobamo enriched near zero and Tobamo enriched near one, plus smaller intermediate mass between zero point four and zero point nine.

**Alt text (Panel f):** Contig-level viral-outside-Virgaviridae counts are concentrated almost entirely below zero point two, with only rare high-probability outliers near one, so most values lie left of zero point five.

| **orf** | **complete or partial [complete + partial]** | **correct predictions** | **false predictions breakdown [tob1,tob2,tob3,oth1,oth2,msa]** | **false prediction rate %** |
|---|---|---|---|---|
| RdRp ORF1 | 59 [3 + 56] | 50 | 9=[0+1+8+0+0+0] | 15.25 |
| RdRp ORF2 | 53 [7 + 46] | 53 | 0=[0+0+0+0+0+0] | 0 |
| MP | 57 [26 + 31] | 53 | 4=[0+3+1+0+0+0] | 7.02 |
| CP | 69 [51 + 18] | 68 | 1=[0+0+1+0+0+0] | 1.45 |

Supplementary Table S3: ORF-level prediction statistics, including breakdown of sequences to complete and partial, number of correct predictions, false-prediction category breakdown, and false prediction rate (%).

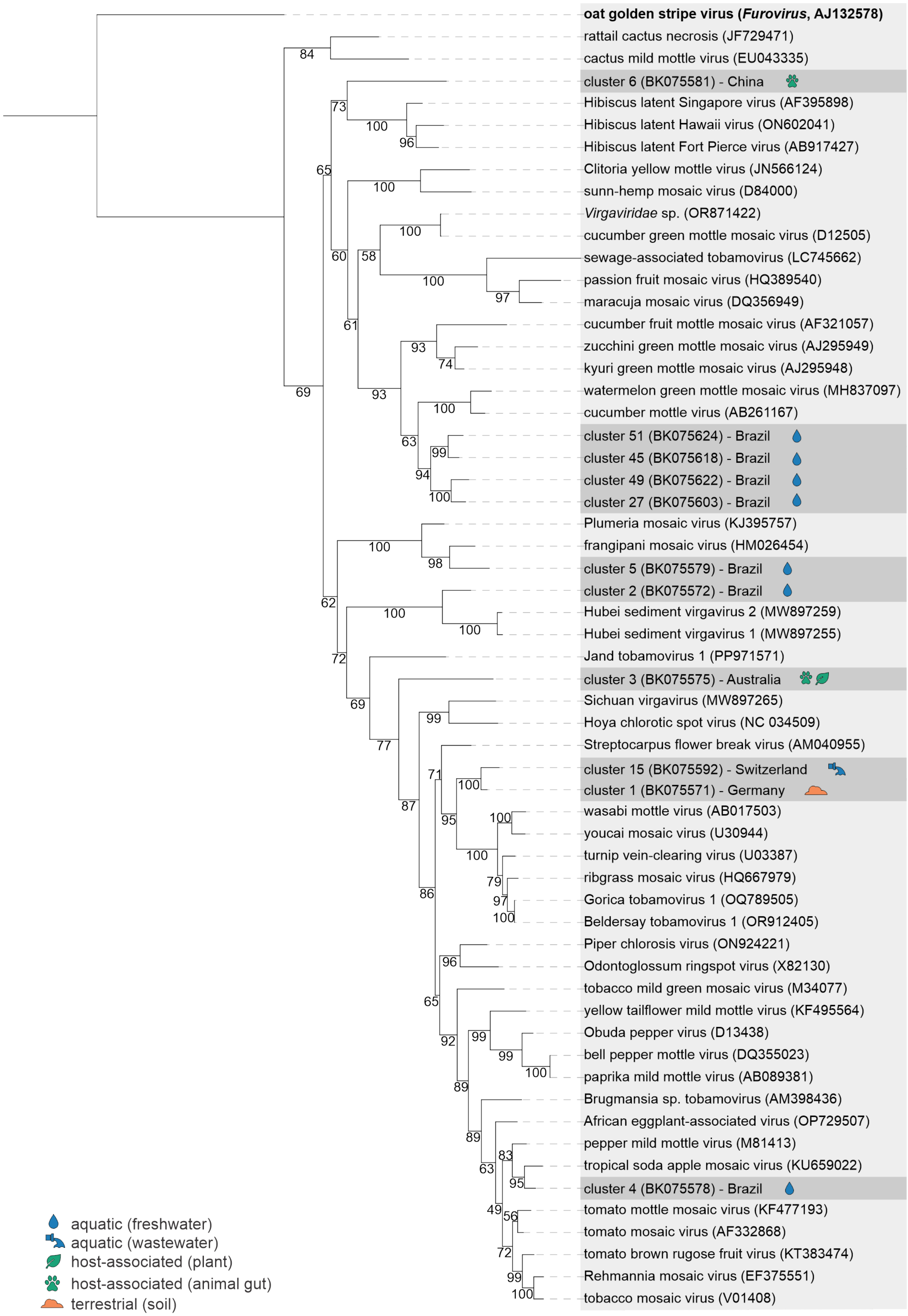

Supplementary Figure S6: (Caption next page.)

Supplementary Figure S6: **Maximum likelihood phylogenetic tree of novel tobamoviral sequences inferred from RdRp ORF1.** The tree is based on the alignment of ORF1 nucleotide sequences from cluster representatives, their closest blastx hits, and the ICTV VMR tobamoviral reference sequences. The tree was inferred in IQ-TREE (v3.0.1) under the TIM2e+G4 model. Support values derived from the ultrafast bootstrap analysis are displayed on the branches. The tree was rooted with an outgroup (oat golden stripe virus, genus *Furovirus*), shown in bold. The cluster representatives are highlighted in dark grey. Icons indicate the source material associated with the datasets used for assembling contigs within each cluster (see legend).

**Alt text:** Maximum likelihood phylogenetic tree constructed from the nucleotide sequences of tobamoviral open reading frame one. The tree was inferred from the alignment of representatives of novel tobamoviruses detected in this study, their closest blastx hits and reference tobamoviral sequences. The tree displays the placement of novel tobamoviral contigs into distinct clades inside the tobamoviral genus.

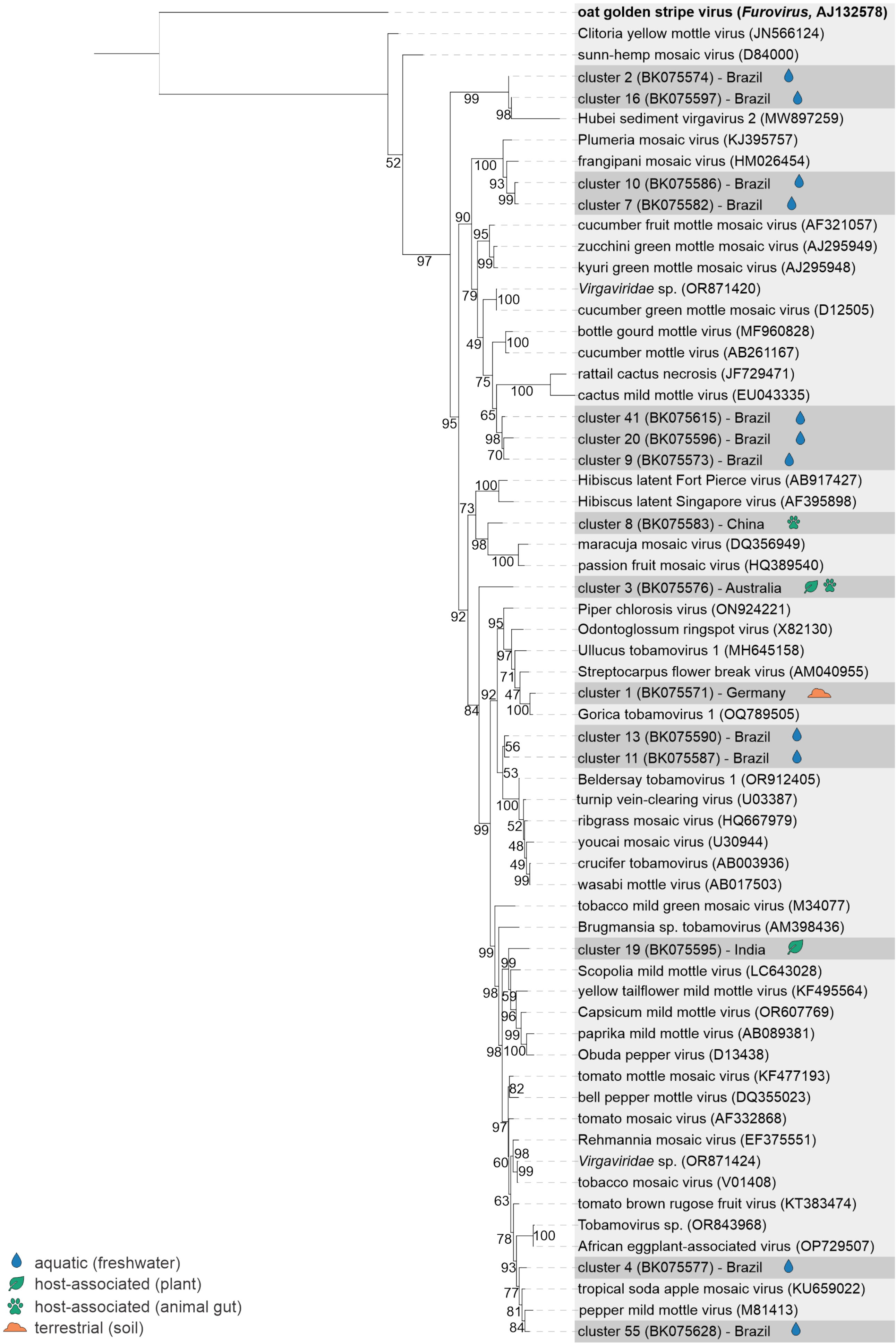

Supplementary Figure S7: (Caption next page.)

Supplementary Figure S7: **Maximum likelihood phylogenetic tree of novel tobamoviral sequences inferred from RdRp ORF2.** The tree is based on the alignment of ORF2 nucleotide sequences from cluster representatives, their closest blastx hits, and the ICTV VMR tobamoviral reference sequences. The tree was infered in IQ-TREE (v3.0.1) under the TIM2e+G4 model. Support values derived from the ultrafast bootstrap analysis are displayed on the branches. The tree was rooted with an outgroup (oat golden stripe virus, genus *Furovirus*), shown in bold. The cluster representatives are highlighted in dark grey. Icons indicate the source material associated with the datasets used for assembling contigs within each cluster (see legend).

**Alt text:** Maximum likelihood phylogenetic tree constructed from the nucleotide sequences of tobamoviral open reading frame two. The tree was inferred from the alignment of representatives of novel tobamoviruses detected in this study, their closest blastx hits and reference tobamoviral sequences. The tree displays the placement of novel tobamoviral contigs into distinct clades inside the tobamoviral genus.

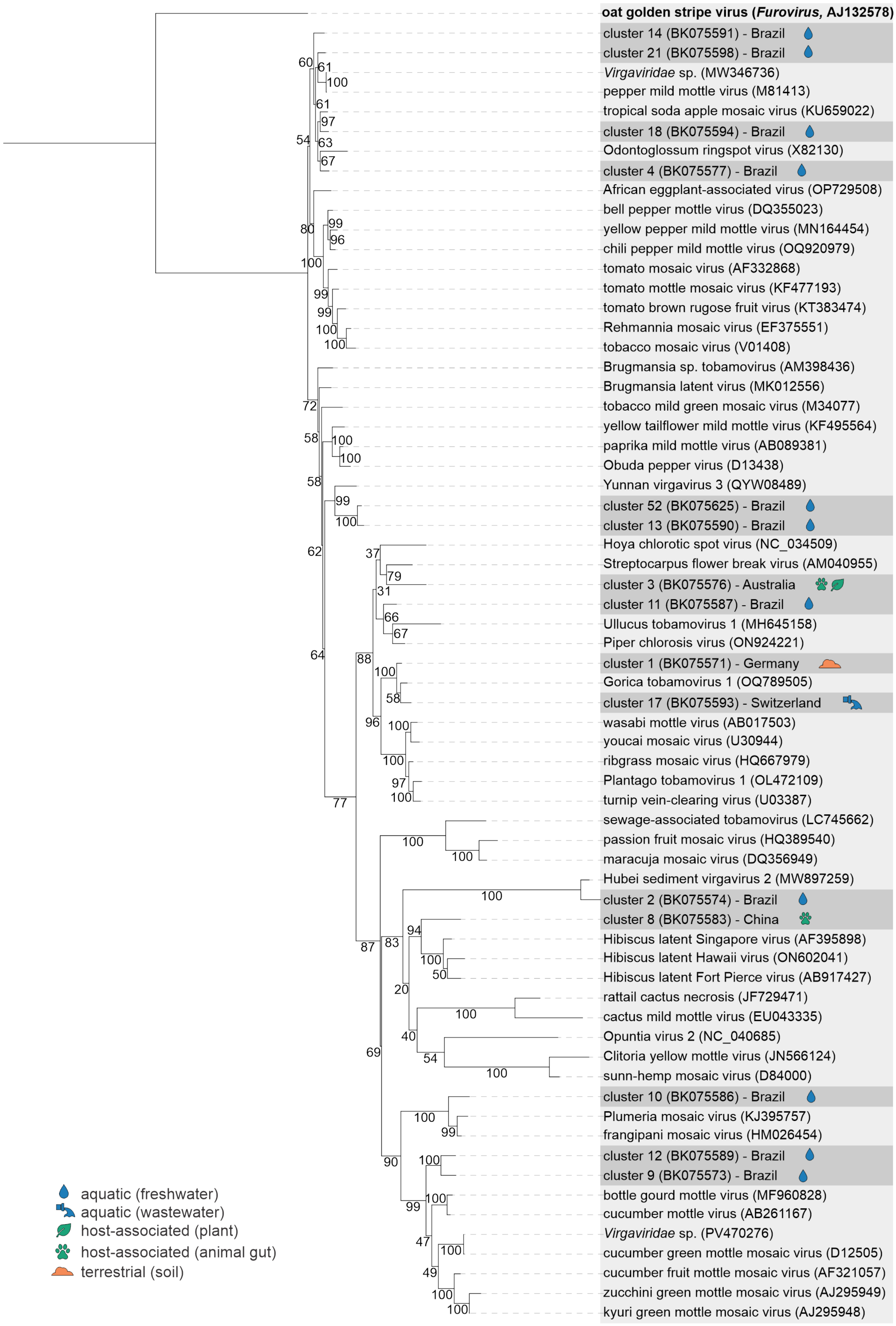

Supplementary Figure S8: (Caption next page.)

Supplementary Figure S8: **Maximum likelihood phylogenetic tree of novel tobamoviral sequences inferred from MP.** The tree is based on the alignment of MP nucleotide sequences from cluster representatives, their closest blastx hits, and the ICTV VMR tobamoviral reference sequences. The tree was infered in IQ-TREE (v3.0.1) under the TIM2e+G4 model. Support values derived from the ultrafast bootstrap analysis are displayed on the branches. The tree was rooted with an outgroup (oat golden stripe virus, genus *Furovirus*), shown in bold. The cluster representatives are highlighted in dark grey. Icons indicate the source material associated with the datasets used for assembling contigs within each cluster (see legend).

**Alt text:** Maximum likelihood phylogenetic tree constructed from the nucleotide sequences of tobamoviral open reading frame three. The tree was inferred from the alignment of representatives of novel tobamoviruses detected in this study, their closest blastx hits, and reference tobamoviral sequences. The tree displays the placement of novel tobamoviral contigs into distinct clades inside the tobamoviral genus.

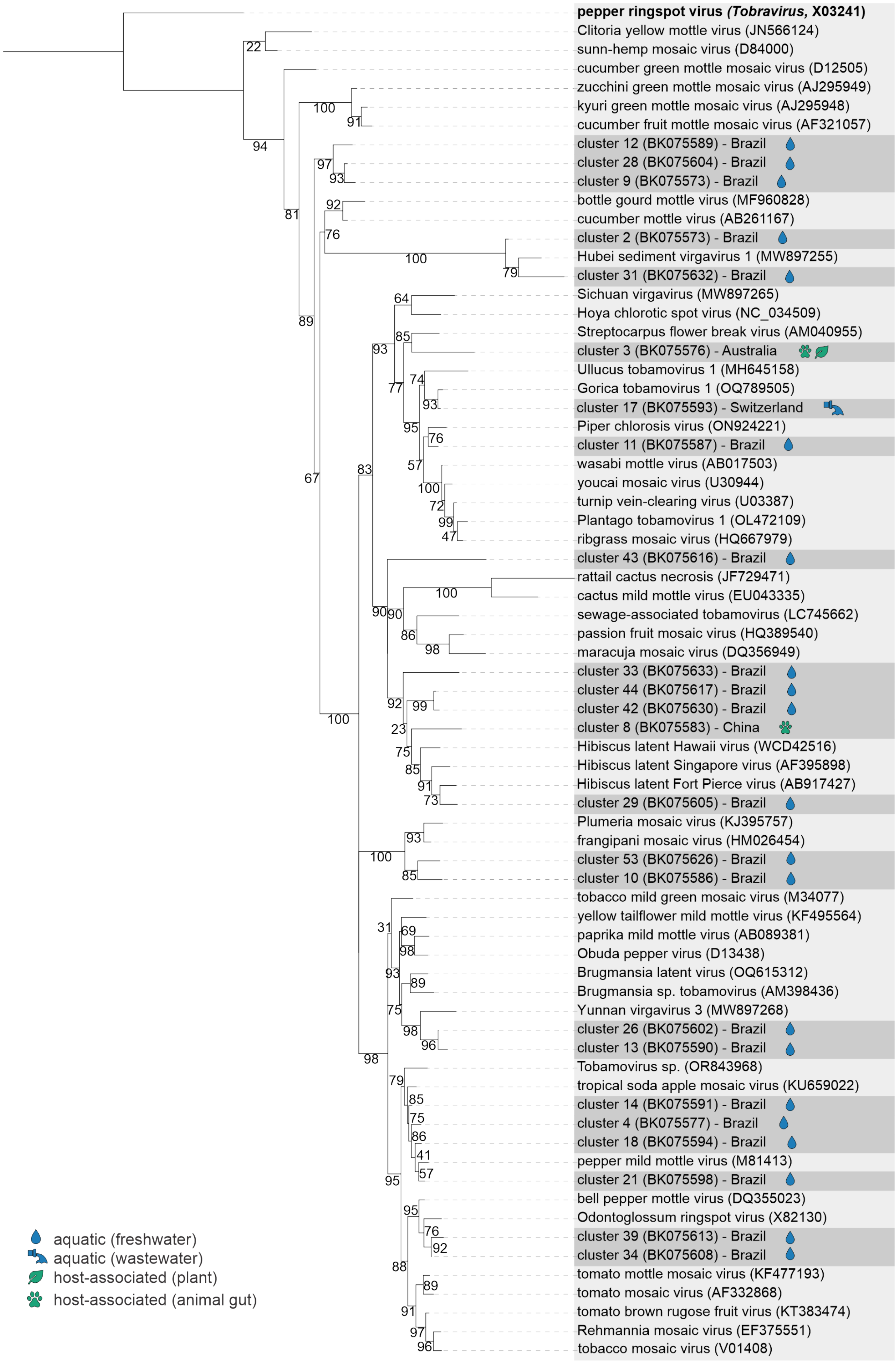

Supplementary Figure S9: (Caption next page.)

Supplementary Figure S9: **Maximum likelihood phylogenetic tree of novel tobamoviral sequences inferred from CP.** The tree is based on the alignment of CP nucleotide sequences from cluster representatives, their closest blastx hits, and the ICTV VMR tobamoviral reference sequences. The tree was infered in IQ-TREE (v3.0.1) under the TIM2e+G4 model. Support values derived from the ultrafast bootstrap analysis are displayed on the branches. The tree was rooted with an outgroup (oat golden stripe virus, genus *Furovirus*), shown in bold. The cluster representatives are highlighted in dark grey. Icons indicate the source material associated with the datasets used for assembling contigs within each cluster (see legend).

**Alt text:** Maximum likelihood phylogenetic tree constructed from the nucleotide sequences of tobamoviral open reading frame four. The tree was inferred from the alignment of representatives of novel tobamoviruses detected in this study, their closest blastx hits, and reference tobamoviral sequences. The tree displays the placement of novel tobamoviral contigs into distinct clades inside the tobamoviral genus.
